# Deep Invaginations of Nuclear Envelope Coordinate Spatial Organization of Chromatin in Epithelium

**DOI:** 10.64898/2026.03.10.710762

**Authors:** Elina Mäntylä, Sanna Korpela, Anna Rekonen, Sini Hakkola, Jenni Karttunen, Anni Pörsti, Sanni Erämies, Fábio Tadeu Arrojo Martins, Roselia Davidsson, Markus J.T. Ojanen, Satu A-M. Hakanen, Peipei Wang, Joonas Uusi-Mäkelä, Alice Anais Varlet, Maija Vihinen-Ranta, Daniel E. Conway, Keijo Viiri, Matti Nykter, Jan Lammerding, Teemu O. Ihalainen

**Author notes:** Corresponding author: Teemu Ihalainen, Address: Faculty of Medicine and Health Technology, Tampere University, Tampere, Finland. Arvo Ylpön katu 34, 33520 Tampere, Finland.

## Abstract

Cell nuclei are often used to assess cell health, but how their shapes vary in normal tissues and how they respond to mechanical forces is not well understood. Here, we describe deep invaginations of the nuclear envelope (DINEs) as common features of epithelial cell nuclei. After their formation, DINEs exist independently of the cytoskeleton, depend on A-type lamins, and emerge in response to cell crowding, contact inhibition, and tissue maturation. High-resolution imaging shows that, in contrast to the peripheral nuclear lamina, DINEs contain densely packed chromatin with regions of active gene transcription. They also remodel dynamically during confined migration, allowing nuclei to adapt to physical constraints. Mechanistically, DINE formation is linked to suppression of MAPK signaling, while activation of growth-promoting pathways reduces their occurrence. These findings reveal DINEs as intrinsic, mechanosensitive structures that coordinate nuclear shape, chromatin organization, and gene activity, providing new insight into how epithelial cells integrate mechanical and biochemical cues to maintain tissue homeostasis.

**Teaser:** Deep nuclear envelope invaginations organize chromatin and gene activity in response to epithelial crowding.

## Introduction

Mechanical forces that arise internally within a cell, from adjacent cells, or from the surrounding microenvironment play critical roles in regulating cellular behavior, and extend their influence to the nucleus (*1*). The nucleus itself is a mechanosensitive organelle that responds to the mechanical cues by undergoing dynamic remodeling, converting changes in cell geometry into downstream signaling that influences cellular functions and behavior (*2*). Alterations in nuclear shape and mechanics co-regulate physiological processes, including differentiation, migration, proliferation, tissue morphology, and homeostasis, and are implicated in the onset and progression of diseases such as cancer (*3–5*). However, how nuclear deformation links to the spatial organization of chromatin and genetic programming remains poorly understood.

Cells sense mechanical forces at the plasma membrane through integrins, cadherins, and other mechanosensitive proteins and their complexes, including focal adhesions and ion channels. These signals are transmitted either mechanically via the actomyosin cytoskeleton or transduced into biochemical signals through the process of mechanotransduction. Mechanical cues are further conveyed to the nuclear envelope (NE) and into the nucleus via the linker of nucleo- and cytoskeleton (LINC) complexes, which physically connect the cytoskeleton to the NE and the underlying nuclear lamina (*5–11*). The nuclear lamina is located beneath the inner nuclear membrane (INM) and is primarily composed of type V intermediate filaments, A- and B-type nuclear lamins (*12*, *13*). The nuclear lamina is a mechanosensitive structure directly responding to mechanical forces exerted on the nucleus (*14*, *15*). In particular, changes in nuclear morphology are accompanied by its structural alterations that affect the accessibility of chromatin-binding lamin epitopes (*15*). The nuclear lamina interacts with chromatin through dynamic, long-range associations involving topologically associating domains and lamina-associated domains (LADs), thereby orchestrating three-dimensional genome architecture and co-regulating gene expression (*16–18*). Mechanical deformation of the nucleus and nuclear lamina can actuate changes in chromatin conformation, epigenetic modifications, and accessibility, ultimately influencing transcriptional activity (*6*, *13*, *19–27*). Importantly, chromatin can directly buffer mechanical stress imposed on the nucleus, thereby protecting genome integrity (*27*). Thus, nuclear lamina-chromatin interactions play a central role in nuclear mechanoadaptation and the orchestration of genome function in response to mechanical stress (*28*).

Over the past decade, it has become evident that NE deformation can be very diverse. Abnormalities in NE morphology, such as dents and blebs, are often associated with nuclear dysfunction and observed in numerous diseases (*29*). Due to these pathological associations, nuclear morphology serves as a diagnostic and prognostic marker, particularly in laminopathies and cancer (*19*, *30*). In contrast, NE invaginations of multiple types are often referred to as the nucleoplasmic reticulum, whose function and physiological relevance remain largely elusive (*31*). Type I invaginations of the INMhave been found to associate with ER markers and nuclear calcium dynamics in fibroblasts and skeletal muscle cells (*31*, *32*). Venturini *et al.* have shown that the INM unfolds upon nuclear stretching, providing physical cues on cell shape distortions and activating calcium-dependent mechanotransduction signaling, which subsequently controls the contractility of actomyosin cytoskeleton and migration plasticity (*2*). Aligned with this, work by Lomakin *et al.* and Eneyedi *et al.* have implicated that NE unfolding and stretching activates calcium-dependent phospholipase cPLA2-mediated contractility, tailoring cell responses to spatial constraints and tissue damage (*33*, *34*). Type II invaginations are double-membrane inward protrusions with a diffusion-accessible cytoplasmic core and have been determined as type II nucleoplasmic reticulum (*31*, *35*). Among these, hollow tube-like invaginations, found in multiple cell types such as fibroblasts and epithelial cells, emanate perpendicular to the basal or apical NE (*31*, *36*). Schoen *et al.* describe the potential role of these structures in the organization of LADs and transcription (*37*). Type II also includes subtle undulations or surface-level wrinkles, which were recently described in breast cancer cells by Wang *et al.* as folding of excess NE surface area, linking this phenotype to increased cellular invasiveness and malignancy (*38*). Relevant to our study, yet distinct type II laterally oriented noncylindrical cytoskeleton-associated deep NE invaginations (which we will abbreviate from here on as DINEs) extending longitudinally across the nucleus have been described (*31*, *39*). These inward NE invaginations that reach deep within the nucleoplasm, traversing the nucleus, and containing cytoskeletal elements within their cytoplasmic core, have been observed in healthy tissues, such as in breast tissue, and in cultured epithelial cells, in both 2D and 3D environments (*40*). However, the role of DINEs has remained unclear. They have been implicated in nucleocytoplasmic communication, gene regulation, calcium signaling in response to mechanical damage, and nuclear lipid metabolism (*13*, *27*, *31*, *41–45*). Additionally, NE wrinkling has been shown to predict the mechano-response of mesenchymal progenitor cells in both 2D and 3D environments (*41*, *46*). In mesenchymal stem cells, NE invaginations have been classified as microtubule-dependent structures that influence formations influencing genome organization by increasing chromatin compaction and reducing accessibility, thereby affecting gene expression (*46*).

In this study, we investigated the nature and function of the DINEs in mature epithelium to reveal how nuclear deformation links to the spatial organization of chromatin and genetic programming. Using both *in vitro* and *in vivo* models, we demonstrate that epithelial nuclei develop DINEs, which traverse the nucleoplasm from the basal to the apical side. Thus, we define DINEs visually as NE invaginations extending longitudinally and inward by at least one half of the apparent nuclear diameter. The DINEs form in conjunction with epithelial maturation, increased cell packing density, and elevated lateral (intercellular) compression, and persist independently of actomyosin or microtubule-mediated cytoskeletal tension. We show that the DINEs are unique nuclear NE deformations: they exhibit basal-like nuclear lamin organization with locally increased accessibility of lamin A epitopes suggesting their formation as protrusions of the basal nuclear membrane and are significantly associated with chromatin. While the chromatin in DINE-containing nuclei is globally compacted with reduced transcriptional accessibility, the DINEs themselves engage differentially with transcriptionally active and accessible chromatin regions, indicating increased chromatin dynamics. Finally, we establish the interdependence between DINEs and reduction of MAPK signaling *in vitro*. We suggest that formation of DINEs is associated with downregulation of MAPK signaling activity. Our study demonstrates that NE invaginations are not merely mechanically induced byproducts but active regulators of spatial chromatin organization and function during the interphase of contact inhibition and EMT-associated phenotypic conversion, thereby introducing a previously unrecognized layer of nuclear regulation.

## Results

### DINEs are prominent nuclear features in mature epithelium both *in vitro* and *in vivo* and exist independently of cytoskeletal organization

During epithelial growth and maturation, cells reorganize themselves to form an intact epithelial layer with barrier functions. We were interested in how nuclear morphology changes during the process *in vitro*. By using laser scanning confocal microscopy (LSCM) imaging of Madine Darby canine kidney type II (MDCK II) cells immunofluorescently labeled for lamin A/C (LA/C) at 7 d post-seeding, i.e., when the monolayer reaches full maturity, we detected that nearly all nuclei (in approximately 85-95 % of nuclei in most experiments of this study) in this confluent monolayer contained wrinkled or invaginated NE (Fig. 1A and Fig. 2A to C, and F). We compared this nuclear phenotype to nuclei in a developing monolayer grown for 2 d and found these nuclei to be mainly smooth, with only ~20 % of them invaginated, which indicated that DINE-containing nuclear phenotype correlated with epithelial maturation (Fig. 1, A and B). Next, we used expansion microscopy (ExM) combined with LSCM to visualize nuclear morphology in super-resolution at 5 min, 2 h, or 7 d post reseeding. For this, we immunostained the cells for lamin B1 (LB1) because of its constitutive and mechanically invariant expression and non-polarized localization at the INM, providing a reliable and stable outline of the nuclear rim. We detected DINEs in the 7 d-grown fully confluent mature monolayers, but these structures were scarce at earlier time points (Fig. 1C). The DINEs were sheet-like NE folds resembling ruffled curtains that extend from the basal side towards apical NE, often dividing the nucleus into spatially distinct lobes (Movie S1). To characterize them further, we visualized the nuclei of 7 d grown cells with transmission electron microscopy (TEM) (Fig. 1D). The TEM confirmed that the DINEs were commonly established at the basal side as inward folding of the NE and could extend close to the apical NE (Fig. 1D). Both LSCM and TEM showed densely packed chromatin and nuclear pore complexes (NPCs) along the DINEs (Fig. 1D and Fig. S1, A to C). Based on these observations, in all experiments of this study we define the DINEs as NE invaginations extending longitudinally and inward visually by at least one half of the apparent nuclear diameter and binding chromatin, which make them unique and distinctive of any other NE undulations. Taken together, these results suggested that DINE-formation is correlated with epithelial growth and maturation. We then confirmed the presence of DINEs also *in vivo* by LA/C immunostaining and LSCM imaging of epithelium-rich mouse esophagus and small intestine (duodenum) tissue sections (Fig. S2), in mouse intestinal organoids, and in human esophagus organoids (Movie S2). Several DINE-containing nuclei were detected within the esophagus mucosa (Fig. S2, A and B), and especially in cells lining the esophagus papillae (Fig. S2C). In the duodenum, a few (1–2) DINE-containing nuclei with pronounced lamin intensity were detected at the bottom of the crypts (Fig. S2D). In human esophagus organoids, DINEs were rare in the interior cell nuclei, but more common in the nuclei of outer-layer cells (Fig. S2E and Movie S2). DINEs were also detected in mouse intestinal organoids but not in Lgr5 positive (Lgr5^+^) intestinal stem cells (Fig. S2F). In line with this, tissue sections from Lgr5^+^ mice intestine showed that whereas Lgr5^+^ stem cells contained smooth nuclei, DINEs were detected in pyramidal/columnar -shaped cells. These DINE-containing cells were located in the crypts in the direct vicinity of Lgr5^+^ stem cells and had cytoplasmic granules and were thus identified as secretory cells (such as Paneth cells) (Fig. S2G), in which irregularly shaped nuclei have been well documented (*47*). This suggests that DINEs are a property of a differentiated epithelial cell *in vivo*. Together, these findings support that DINEs are not merely an artifact arising in *in vitro* culture of an epithelial monolayer but that DINEs also occur in epithelial 3D organoids and epithelial tissues *in vivo*.

**Fig 1.**
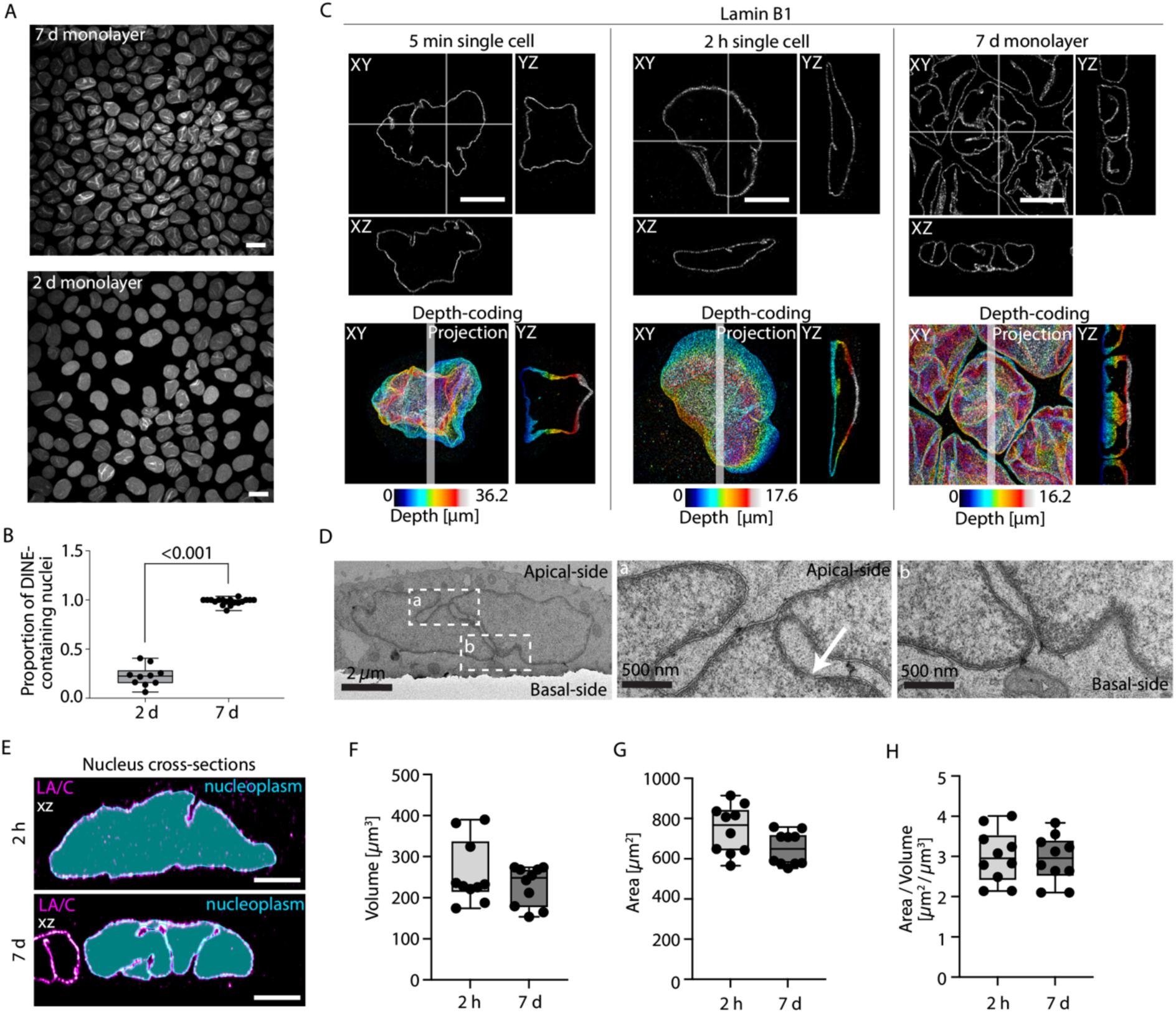
Deep invaginations (DINEs) of the nuclear envelope are prevalent features of epithelial nuclear architecture. **(A)** Confocal fluorescence microscopy image of LA/C-rod -stained epithelial cells showing nuclear morphology and emergence of DINEs *in vitro*. **(B)** Box-and-whiskers plot presenting quantification of the proportion of DINE-containing nuclei at 2 d and 7 d grown epithelium, n= three independent biological**(C)** Expansion microscopy images and their depth-color coded surface renderings displaying NE morphology and DINEs at 5 min, 2 h and 7 d post-seeding. Scale bars, 5 µm. **(D)** Transmission electron microscopy presenting DINE morphology in a single epithelial nucleus at 7 d. Scale bars, 2 µm (upper panel) and 500 nm (middle and lower panels). **(E)** Representative segmentations of NE and nucleoplasmic regions of nuclei at 2 h and 7 d post-seeding by using lamin A/C (LA/C) immunostaining. Box-and-whiskers plots showing quantification of **(F)** nuclear volume, **(G)** total NE surface area (all NE surface containing DINEs), and **(H)** NE total area-to-volume -ratio from single nuclei. In A-G, n=three independent biological replicates. Box-and-whisker plots represent the 25^th^–75^th^ percentiles as boxes, the median as a line within the box, and whiskers indicating the minimum and maximum values.

**Fig 2.**
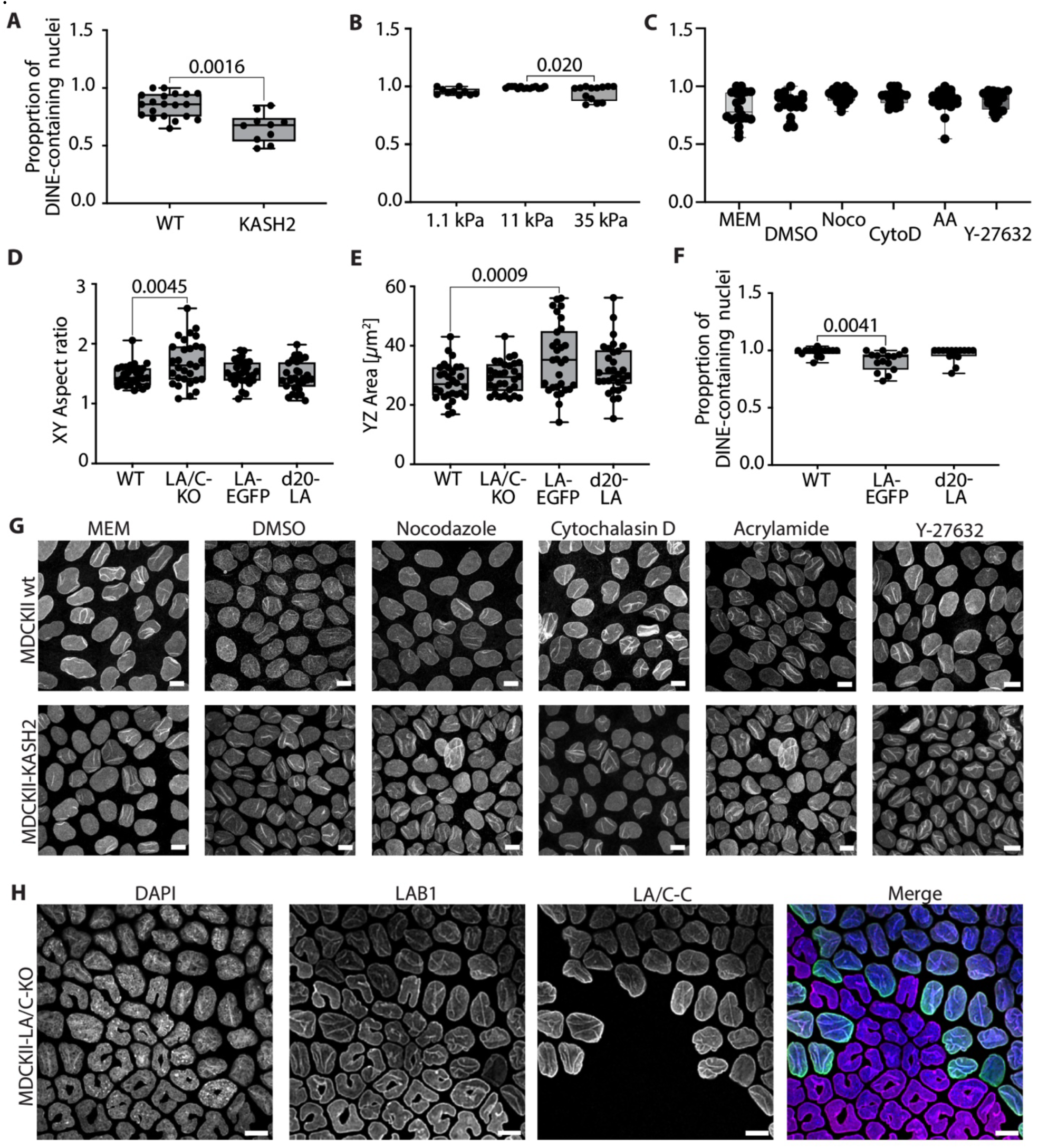
DINEs are insensitive to actomyosin-generated tension and extracellular matrix stiffness but are contingent on lamin A/C. Box-and-whiskers plots showing proportions of DINE-containing nuclei in **(A)** wild-type (wt) (n=25 fields, three independent biological replicates) and KASH2-dominant negative (n=15 fields, two independent biological replicates, unpaired two-tailed Student’s T-test), **(B)** 1.1, 11 and 35 kPa surface rigidities (n=10, 14 and 13 fields, three independent biological replicates), and **(C)** complete growth medium (MEM), dimethyl sulfoxide (DMSO, control), or cytoskeletal drug -treated (nocodazole, cytochalasin D, acrylamide, and ROCK-inhibitor Y-27632) treated cells at 7 d post-seeding (n=20 fields, three independent biological replicates). Box-and-whiskers plots indicating nuclear **(D)** XY aspect ratio (n=60, 60, and 30 cells, three independent biological replicates), **(E)** YZ area, (n=30 cells in each, three independent biological replicates) and **(F)** proportion of DINE-containing nuclei in wt, lamin A/C knock-out (LA/C-KO), lamin A-EGFP (LA-EGFP) and lamin A delta20 (N-terminal deletion of 20 amino acids; d20-LA) expressing cells (n=15 fields, three biological replicates, One-way ANOVA with Dunnett’s multiple comparison test, p<0.05 indicates statistical significance). Box-and-whisker plots represent the 25^th^–75^th^ percentiles as boxes, the median as a line within the box, and whiskers indicating the minimum and maximum values. **(G)** Confocal microscopy images showing nuclear morphology in lamin A/C-rod immunostained 7 d-grown epithelial cell nuclei following treatment with cytoskeletal drugs shown in (C). **(H)** Confocal microscopy image showing absence of DINEs in lamin A/C-deficient (LA/C-KO) nuclei in comparison to wt nuclei following LA/C-C-terminus (LA/C-C) immunostaining. Scale bars, 10 µm. In B-F, One-way ANOVA, Dunnett’s multiple comparison test, p<0.05 indicates statistical significance.

We found that the DINEs divided the nucleus into nearly spatially distinct lobes, where the basal NE almost traversed through the nucleus. We speculated that DINEs could affect the nuclear surface-to-volume ratio. We addressed this by quantifying the nuclear volume, the total NE surface area including all peripheral NE and the DINEs, and the surface area-to-volume -ratio in mature cells grown for 7 d and in cells grown for 7 d and reseeded for 2 h, showing smooth or DINE-containing nuclei, respectively. We manually segmented the NE from ExM-LSCM z-stacks based on LA/C staining (Fig. 1E). The average nuclear volume (mean ± standard deviation, SD) had no statistically significant differences at 7d and 2 h post-seeding (230 ± 50 µm³ and 260 ± 80 µm³, respectively) (Fig. 1F). Similarly, the total NE surface area, was highly similar at 7 d (650 ± 90 µm²) and 2 h (750 ± 120 µm²) post-seeding (Fig. 1G). The total surface area-to-volume ratio was also comparable between the two time points (2.9 ± 0.6 at 7 d and 3.0 ± 0.7 at 2 h) (Fig.1H). Since the folded DINEs accounted for a large fraction of the NE at 7 d post-seeding, the easily accessible NE area appeared reduced at this time point. Overall, the data suggest that DINEs form through remodeling of the existing NE rather than expansion of its total surface area.

We then sought to test how the DINEs are formed and maintained. The role of cytoskeleton, especially actin, in regulating nuclear morphology is well documented (*43*, *48*, *49*). Nuclear invaginations in several cell types have been linked to mechanical forces generated or transmitted by actin, microtubules, or intermediate filaments (*42*, *43*, *50–54*). Therefore, we asked whether DINEs depend on functional LINC complexes, actomyosin cytoskeleton, substrate rigidity, or cellular contractility. We investigated the role of LINC-complexes by determining the proportion of DINE-containing nuclei in WT and cells stably expressing the dominant negative KASH domain of Nesprin-2 (EGFP-KASH2), which disrupts the LINC complexes and thus weakens the nucleo-cytoskeletal coupling (*55*). The analysis at 7 d indicated that disruption of the LINC complexes slightly reduced the proportion of DINE-containing nuclei. While ~85 % of WT cells contained DINEs, ~66 % of EGFP-KASH2 cells displayed nuclear folding, suggesting that DINEs are affected by but not completely dependent on LINC coupling (Fig. 2A). We then tested the effect of substrate rigidity, a known modulator of cellular contractility, on the existence of the DINEs. MDCK II WT cells were cultured for 7 d on polyacrylamide (PAA) hydrogels of three rigidities (Young’s modulus of 1.1, 11, and 35 kPa) (Fig. 2B). The proportion of DINE-containing nuclei was high on all growth surface rigidities (~96 %, ~99 %, ~95 % in 1.1, 11, and 35 kPa gels, respectively). We also investigated DINE maintenance and nuclear morphology in WT (Fig. 2, C and G, and Fig. S1, D to F) and EGFP-KASH2 (Fig. S1, G to I) expressing cells grown for 7 d followed by a 2 h incubation with drugs depolymerizing actin (Cytochalasin D), microtubules (nocodazole), or intermediate filaments (acrylamide, AA) (see also Table S1). In addition, DINEs and nuclear morphology were quantified following reduction of cellular contractility with inhibition (2 h) of the ROCK pathway (Y27632). In these studies, dimethylsulfoxide (DMSO) was used as a negative control. None of the treatments abolished DINEs in WT (Fig. 2C) or EGFP-KASH cells (Fig. S1G), indicating that actomyosin cytoskeleton and/or contractility are not required to maintain the DINEs at this timescale. However, these results do not rule out the possibility that cytoskeletal components may be required for the initial DINE formation.

A-type lamins are key regulators of nuclear mechanics, influencing nuclear stiffness and shape (*56*, *57*). We therefore investigated the role of the A-type lamins in DINE formation by quantifying the DINEs and nuclear morphology in *LMNA* gene knockout (LA/C KO) cells (*58*), in stably lamin A-overexpressing (LA-EGFP) cells, and in cells stably expressing polymerization-deficient LA mutant lacking the first 20 amino acids (d20-LA-EGFP). After 7 d in culture, the cells were fixed and analyzed for their nuclear XY aspect ratio (AR) and YZ cross-section area, and the proportion of DINE-containing nuclei in the population. LA-EGFP and d20-LA-EGFP nuclei appeared to have nuclear morphology similar to WT nuclei, but LA/C KO nuclei were highly irregular, with deformed and curved nucleus shape. This was seen in the nuclear XY ARs between the studied cell lines (mean AR ± SD); 1.5 ± 0.2 in WT; 1.5 ± 0.2 in LA-EGFP; 1.5 ± 0.2 in d20-LA-EGFP; and1.7 ± 0.4 in LA/C KO) (Fig. 2, D and H). The YZ cross-sectional nuclear area was significantly larger in the LA-EGFP (36.0 ± 12 µm^2^) in comparison to that of the WT (28 ± 6.1 µm^2^). The YZ nuclear areas in d20-LA-EGFP (32 ± 8.6 µm^2^) and LA/C-KO (29 ± 5.0 µm^2^) were highly similar to that of the WT (Fig. 2E). In WT, ~98 % of the nuclei contained DINEs (Fig. 2F). The proportion of DINE-containing nuclei in the d20-LA-EGFP was highly similar (~97 %), but slightly lower in LA-EGFP expressing cells (~91 %) (Fig. 2F). Intriguingly, while irregularly shaped LA/C KO nuclei exhibited various NE anomalies like dents and grooves, DINEs comparable to those in WT were not observed (Fig. 2H). These results indicated that lamin A levels and organization influence nuclear shape and DINE prevalence. Overexpression of LA slightly increased the nuclear cross-sectional area and reduced DINE frequency, whereas in LA/C KO cells DINEs were not detected and the nuclei were abnormal in shape, showing that LA/C is required for nucleus shape regulation and DINE formation.

Together, our results show that DINEs are prevalent components of the epithelial nuclear architecture both *in vitro* and *in vivo.* DINEs contain both A- and B-type lamins as well as NPCs, which suggests that they represent functional regions of the NE. Like nuclear morphology, the formation of DINEs appears to depend on A-type lamins and occurs independently of growth surface rigidity. Once established, DINEs are stably maintained independently of cytoskeleton organization or force generation.

### DINE formation correlates positively with increasing cell packing density and DINEs undergoes dynamic remodeling during wound healing and confined 3D migration

As DINEs were detected in the fully confluent 7 d old mature epithelium, we thus studied further the influence of growth density, cellular packing, and epithelial maturation on nuclear morphology by introducing growth restriction by surface patterning. Collagen coated hourglass-shaped adhesive islands (400 µm long, 100 µm wide, 10 µm at the narrowest point, the ‘waist’) were fabricated on coverslips to allow the generation of a gradually increasing cell density array on the pattern (Fig. 3A). Occluding-emerald expressing MDCK II cells were grown on the patterns for 3 d until intact monolayer was formed (Fig. 3B), or for 7 d until full confluency and maturity (Fig. S3), fixed, and stained with actin phalloidin and a LA/C antibody. Cells grown for 3 d formed a monolayer with the highest cell density at the waist, and at the vertices of the pattern (Fig. 3C). Cell growth and shape at the waist and edges of the pattern were restricted by the growth surface. Based on qualitative visual assessment at 3 d, nuclei in the waist and edge regions of the pattern corresponding to areas of highest cell density appeared highly wrinkled, although pronounced invaginations were infrequent. In contrast, nuclei in the more sparsely populated central regions of the patterns appeared smoother and more rounded. (Fig. 3D). In comparison, growing cells for 7 d resulted in a high cell density with DINE-containing nuclei throughout the pattern (Fig. S3). These results indicate that epithelial nuclei begin to deform in response to increasing cellular packing density during restricted growth, but the formation of the DINEs correlates with increased cellular packing density and epithelial maturation.

**Fig. 3.**
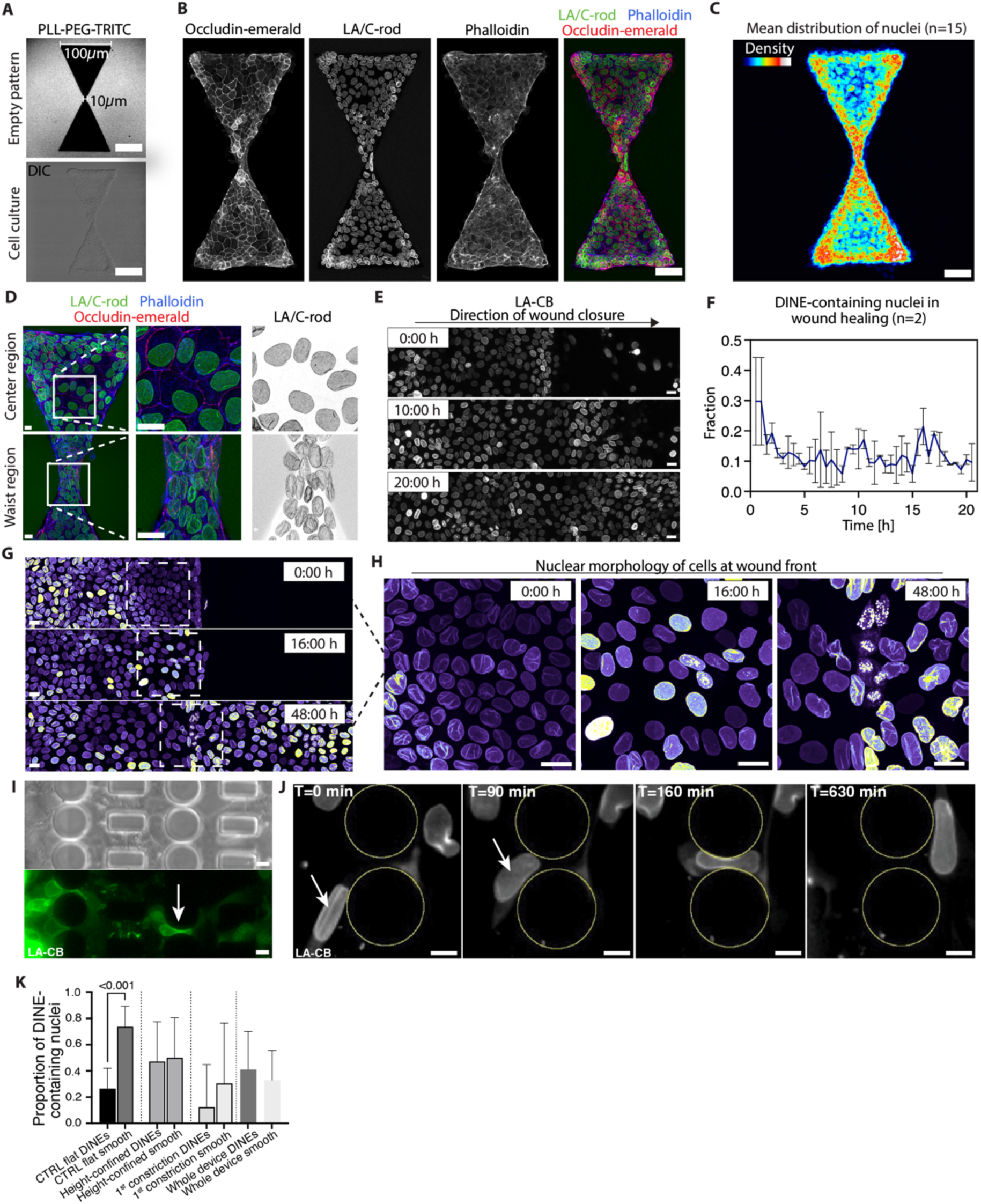
Dynamics of DINEs are affected by spatial confinement of growth environment. **(A)** PLL-PEG-TRITC-stained hourglass-shaped cell adhesive islands (400 µm long and 100 µm wide with 10 µm waist) without (upper panel) and with cells (difference interference contrast (DIC), lower panel), n=three independent biological replicates. Lamin A/C-rod domain (LA/C-rod) immunostained Occludin-Emerald - expressing MDCK II cells **(B)** grown on adhesive island for 3 d, showing DINE-containing nuclei at the pattern vertexes and the waist with increased cellular packing density, and **(C)** mean density distribution of nuclei from n=15 patterns, three independent biological replicates. Scale bars, 50 µm, n=three independent biological replicates. **(D)** Zoom-in images of the pattern center and waist regions. Scale bars, 10 µm. **(E)** Wound healing live cell LSCM of MDCK II cells expressing EGFP-tagged chromobody targeting A-type lamins (LA-CB) - indicating distribution of cells and nuclear morphology at 0, 10, and 20 h post-wounding. Scale bars, 20 µm. **(F)** Quantification of the fraction of DINE-containing nuclei in the wound between 0-20 h post-wounding. Error bars present standard deviation (SD), n=two independent biological replicates. **(G)** Pseudo-colored LSCM images at fixed time points (0, 16, and 48 h) post-wounding, illustrating nuclear morphology at the wound front. Signal intensity is displayed using a pseudo-color lookup table to visualize relative intensity differences. **(H)** Higher-magnification views of the corresponding time points. Scale bars, 20 µm. **(I)** Live cell imaging in a microfluidic 3D confined migration device (upper panel) with LA-CB cells (lower panel), n=seven independent biological replicates. **(J)** Representative single LA-CB cell image indicating unfolding of a DINE during 3D migration through a 2 × 5 µm^2^ (height × width) constriction and long after the migration, 6 fph. Scale bars, 10 µm. **(K)** Quantification showing proportion of DINE-containing LA-CB nuclei (%) in cells grown for 4 d on glass (CTRL), cells migrated into the confinement entry area, cells within the 1st constrictions in 3D microfluidic devices, and cells within the whole devices. Bar plots represent mean ± SD. One-Way ANOVA with Dunnett’s multiple comparisons test, p<0.05 indicates statistical significance.

We then examined how cellular growth density and cell packing dynamics over time influence nuclear morphology. For this, we observed DINE-containing nuclei during collective 2D migration in wound healing, where changes in cell density are coordinated at the population level and linked to epithelial-to-mesenchymal transition (EMT) signaling. For this, MDCK II cells stably expressing EGFP-tagged chromobody against A-type lamins (LA-CB) were grown for 4 days on glass coverslips with a polydimethylsiloxane (PDMS) stencil. Following the creation of a wound by removal of the stencil (at time = 0 h), the cells were allowed to migrate into the wound for 20 h to observe collective migration at the wound front. In this assay, the same field of view with the wound was imaged at 30 min intervals (Fig. 3E). From the timelapse, the ratio of DINE-containing nuclei and the total number of nuclei within the field of view was determined over time. From the stencil liftoff at timepoint 0 h, the fraction of DINE-containing nuclei within the population first declined significantly and remained constant during the collective movement (Fig. 3, E and F). The population of cells at the wound front showed DINEs at 0 h, lost them during collective migration, and regained them after the wound closure, especially following re-maturation of the epithelium (48 h) (Fig. 3, G and H). These dynamic changes in nuclear morphology indicate that EMT-related processes influence existing DINEs and may drive their unfolding.

Next, we wanted to understand if DINEs were associated with nuclear deformability in a highly confined environment and to study their dynamics during constricted 3D migration. We performed live cell imaging of the confluent monolayer of LA-CB cells grown for 4 days in microfluidic 3D migration devices that mimic confined interstitial spaces(*59*) and followed their nuclear morphology with time lapse imaging with 10 min intervals for 24 h during 3D migration along an essential growth factor gradient established with fetal bovine serum (FBS, from 2 % to 10 %) (Fig. 3I and Fig. S3C). The cells were initially seeded into ~200 µm-tall reservoirs, which are not 3D confined spaces. From these reservoirs, the FBS gradient sparked migration into the devices. In the devices, cells first moved within a 3D height-confined entry region located in front of the constrictions and then entered the constriction area, which contained subsequent rows of narrow 2 × 5 mm^2^ constrictions that require significant nuclear deformation during the migration (*59*, *60*). As a control, cells were grown for 4 d in a non-confined environment on glass. While migrative behavior is not characteristic for epithelial cells in intact tissues, and these cells also tended to be less migratory within our 3D migration devices, this experiment enabled us to explore DINE dynamics during non-collective, constricted movement and to determine whether the DINEs require cell-cell interaction. In comparison to control cells grown on glass, in which ~26 % exhibited DINEs (mean proportion of DINE-containing cells within population), around 47 % of the cells within the confined entry area contained DINEs, showing that the height-confined microenvironment and presumably higher cell packing density promoted the formation of DINEs (Fig. S3C). In contrast, on average, only ~12 % of cells that had entered the first row of constrictions contained DINEs, which indicates that either cells without DINEs were entering the constrictions or that DINEs became unfolded before the nucleus squeezed through a constriction (Fig. 3J and Movie S3). Some cells also formed DINEs within the device, as ~38 % of all cells located within the device between the first and the last constrictions contained DINEs. The data shows that the DINEs can be dynamically remodeled during 3D migration. Together, these results support the view that DINEs are dynamic structures that form and unfold in response to changes in cellular packing density and physical confinement.

### DINEs exhibit a basal-like nuclear lamina organization characterized by locally increased accessibility of lamin A

The accessibility of LA/C C-terminal part and structural organization of lamins change in response to physical modulation of the nucleus (*15*, *61*, *62*). To study the structural organization of the nuclear lamina in the DINEs, we used ExM and ratiometric LSCM imaging, in which increased LA/C-C intensity suggests increased antibody binding and thus increased epitope accessibility (Fig. 4A-B). The fluorescence intensities at the apical and basal side of the nucleus were measured from a line drawn in deconvoluted and orthogonally resliced images along a feature, and intensities at the DINEs were measured from xy-images and divided by the factor of 2 correcting for doubled nuclear lamina in the DINE-region following immunostaining against the LA/C-rod, LA/C-C and LB1. The mean fluorescence intensity of LA/C-rod was equal between the apical (mean ± SD, 26.0 ± 9.0 arbitrary units (a.u.)) and basal (27.0 ± 7.0 a.u.) sides of the nucleus (Fig. 4C). However, LB1 was polarized to the apical side (23.0 ± 5.0 a.u.) in comparison to that on the basal side (18.0 ± 4.0 a.u.) (Fig. 4D). Within the DINEs the LB1 (16.0 ± 4.0 a.u.), LA/C-rod (25.0 ± 5.0 a.u.) and LA/C-C (11.0 ± 3.0 in LB1-stainings and 20.0 ± 3.0 a.u. in the LA/C-rod stainings) intensities were found to be similar to those observed on the basal side (Fig. 4C-D). The analysis also indicated that the fluorescence intensity ratio (describing the nuclear lamina composition, mean ± SD) LA/C-C : LA/C-rod was similar across the DINE (0.5 ± 0.2), basal (0.6 ± 0.2), and apical sides (0.4 ± 0.2) (Fig. 4C). In comparison, the LA/C-C : LB1 -ratio was similar between DINEs (0.7 ± 0.2) and the basal side (0.6 ± 0.2), but both were higher than at the apical side (0.3 ± 0.2) (Fig. 4D). The results indicated that in comparison to the apical side, the accessibility of LA/C was similarly increased within the DINEs and at the basal side. When considering our results from the surface area-to-volume analyses described in Fig. 1D-G, these findings support the interpretation that DINEs originate as basal NE inward protrusions.

**Fig. 4.**
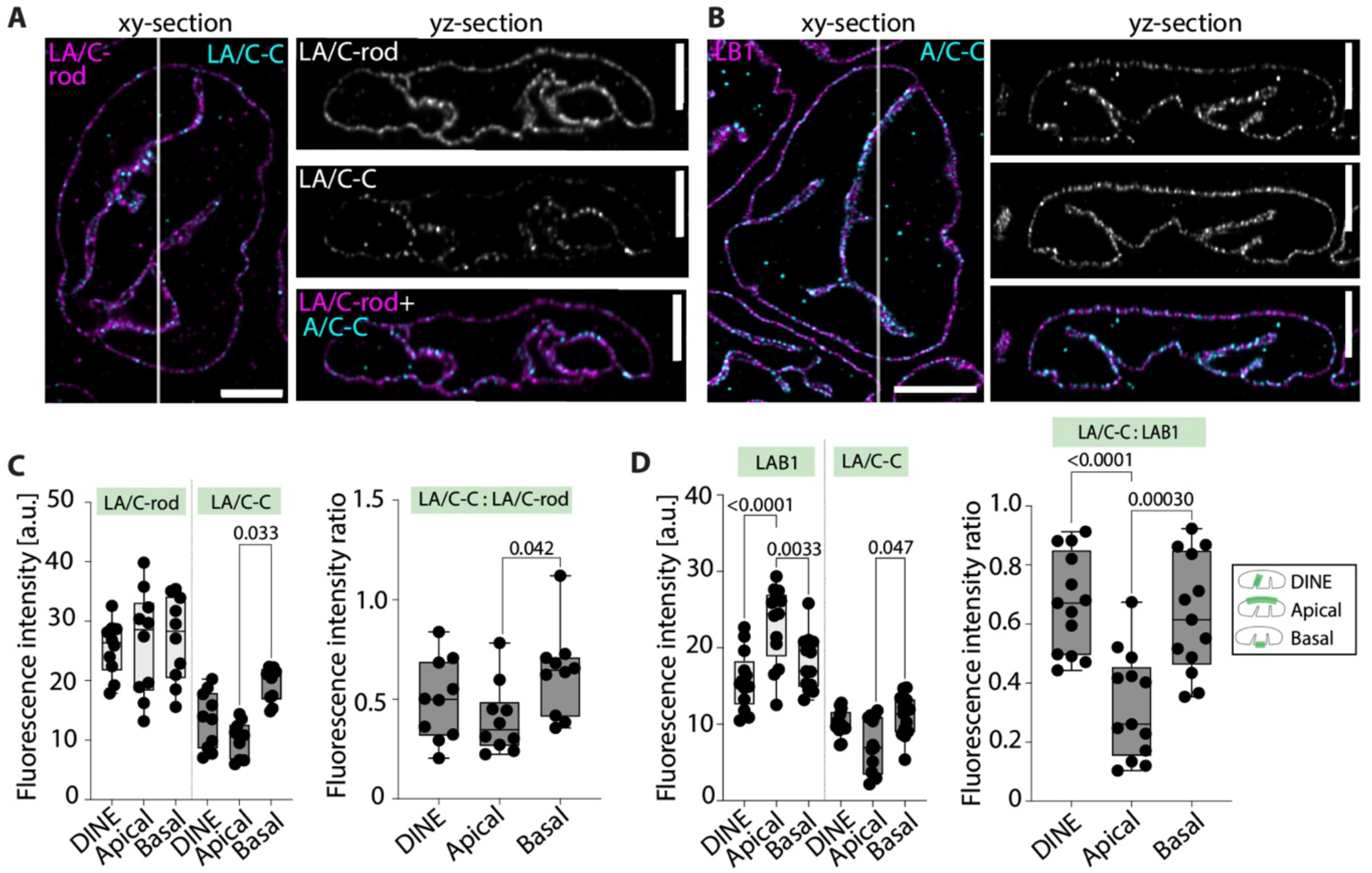
DINEs contain a basal-like nuclear lamina organization. Representative xy- and yz -section expansion microscopy (ExM-LSCM) images from **(A)** LA/C-rod and LA/C-C (two left panels) and **(B)** LB1 and LA/C-C (two right panels) immunostained MDCK II nuclei showing distribution of lamin intensities within the DINEs and on the apical and basal NE. Scale bars, 5 µm. **(C)** and **(D)** Box-and-whisker plots showing quantifications of LSCM-derived lamin intensities (left) and their intensity ratios (right) within the DINEs or on the apical and basal NE. n=three independent biological replicates. One-way ANOVA with Dunnett’s multiple comparisons test indicates statistical significances. Box-and-whisker plots represent the 25^th^–75^th^ percentiles as boxes, the median as a line within the box, and whiskers indicating the minimum and maximum values.

### DINEs contribute to the spatial and functional organization of chromatin

The C-terminal region of LA/C (recognized by the LA/C-C antibody used here) is known to mediate direct chromatin binding (*51*). Changes in the organization of the nuclear lamina and the accessibility of this C-terminal region have been proposed to influence the binding, organization, and gene expression of lamin-interacting genomic regions (*63*). We thus sought to analyze chromatin organization and transcriptional status within the DINEs and elsewhere in the nucleus. The TEM imaging (Fig. 1C) showed that both the DINEs and the peripheral nuclear rim were associated with an electron-dense layer, indicative of densely packed chromatin. To further analyze the chromatin in the vicinity of the lamina, we used ATAC-see (*64*), which is an imaging-based adaptation of the ATAC-seq technique that uses a hyperactive Tn5 transposase loaded with fluorescently labeled adapters to bind and tag open and transcriptionally accessible chromatin *in situ,* enabling direct visualization of open chromatin regions (Fig. 5A and 5B). ATAC-see-tagged accessible chromatin and the total DNA detected with DAPI (4′,6-diamidino-2-phenylindole), were quantified within 0-600 nm distance from the DINE or along the NE rim (referred hereafter as the edge) (Fig. 5A-5C). For each fluorescence profile, we normalized the DAPI and ATAC-see intensities to their local maxima in the 0-600 nm range. Next, we calculated the ATAC-see : DAPI intensity ratio to estimate the fraction of accessible chromatin close to the lamina. DINEs had more chromatin near the lamina (higher DAPI intensity) than the nuclear edge (Fig. 5C). In the immediate vicinity of the lamina (<40 nm), the ATAC-see : DAPI -ratio was lower at DINEs (0.86 ± 0.13) than at the edge (1.75 ± 1.21), indicating a smaller fraction of accessible chromatin at DINEs (Fig. 5C). To further examine the transcriptional status of the chromatin at the DINEs, we labelled DNA with DAPI and immunostained active, serine 2 (S2) phosphorylated RNA polymerase II (pRNApolII) (Fig. 5D). Normalized pRNApolII intensity profiles showed increased intensity at the immediate vicinity (at 0-40 nm distance) of the DINEs (0.64 ± 0.22) compared to nuclear edge (0.19 ± 0.11) (Fig. 5E). Similarly, the pRNApolII:DAPI intensity ratio (<40 nm) was higher at DINEs (0.83 ± 0.31) than at the edges (0.46 ± 0.22) (Fig. 5E). Notably, these ratios were lower than those measured deeper in the nucleus, where transcriptionally active chromatin is known to be located. Taken together, these results suggest that DINEs have more chromatin near the lamina than the nuclear edge. Furthermore, enrichment of active RNApolII at DINEs suggests that the invaginations may influence spatial chromatin organization and transcriptional activity.

**Fig. 5.**
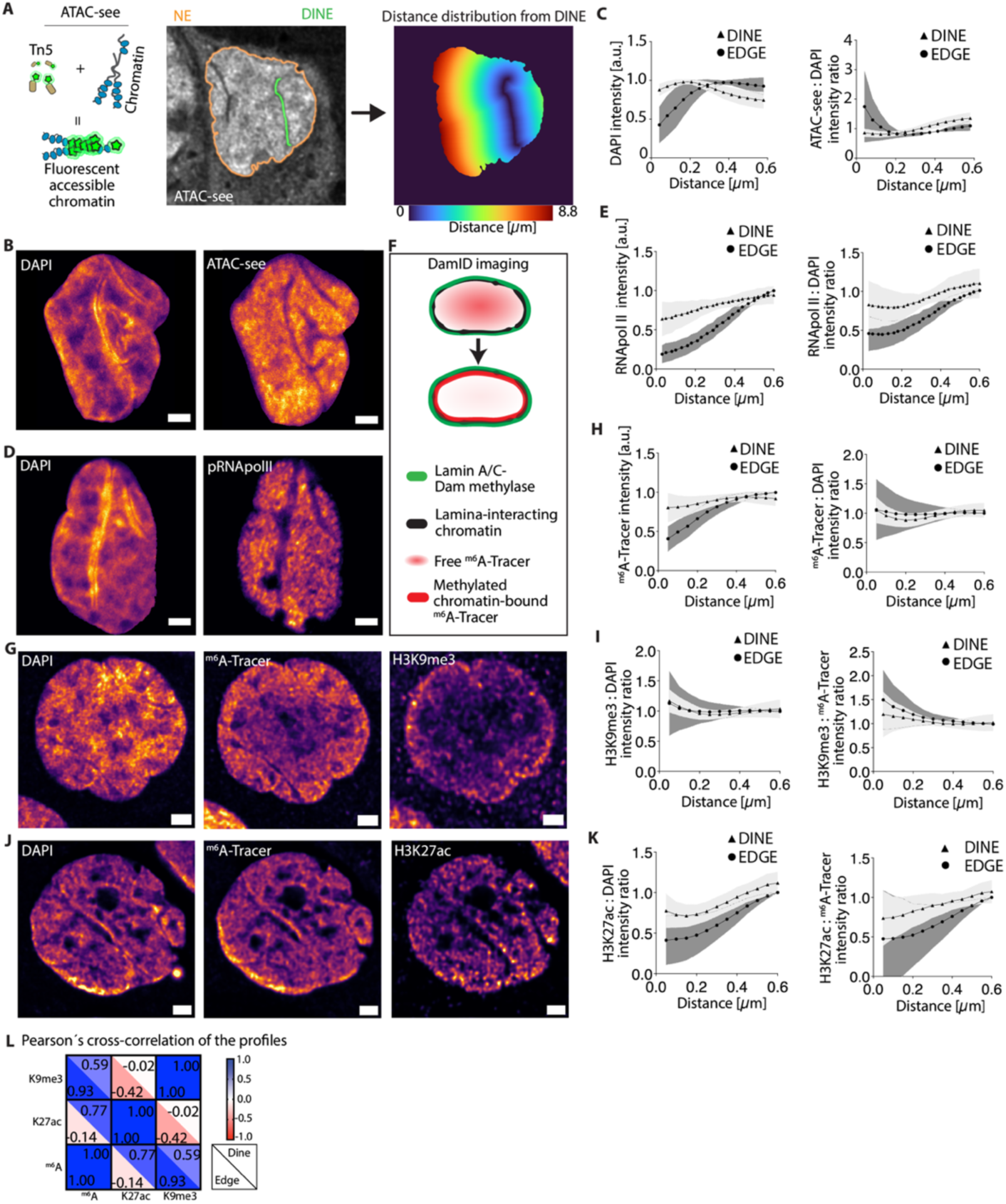
DINEs contribute to spatial and functional 3D chromatin organization. **(A)** Schematic presentation of ATAC-see (left), LSCM image of a single epithelial nucleus after ATAC-see showing peripheral NE (edge, orange) and a DINE (green) (middle), and Euclidian distance analysis map with colors indicating distance from a DINE (µm) (right). **(B)** LSCM images representing intranuclear distribution of heterochromatin (DAPI, left) and transcriptionally accessible chromatin (ATAC-see, right). **(C)** Mean Euclidian distance (0-0.60 µm) distribution from the DINE of DAPI (left) and ATAC-see : DAPI -ratio (right). **(D)** LSCM images representing intranuclear distribution of DAPI (left) and active phosphorylated RNApolII (pRNApolII, right). **(E)** Mean Euclidian distance distribution from the DINE of pRNApolII (left) and pRNApolII : DAPI -ratio (right). **(F)** Schematic presentation of LA/C-Dam ID showing redistribution of LAD marker ^m6^A-Tracer-mScarlet following induction of the lamin-conjugated methylase. **(G)** LSCM images representing intranuclear distribution of DAPI (left), LADs ^(m6^A-Tracer, middle) and transcriptionally repressed chromatin (H3K9me3, right). **(H)** Mean Euclidian distance distribution from the DINE of ^m6^A-Tracer (left) and ^m6^A-Tracer : DAPI -ratio (right). **(I)** Mean Euclidian distance distribution from the DINE of H3K9me3 : DAPI- (left) and H3K9me3 : ^m6^A-Tracer -ratios (right). **(J)** LSCM images representing intranuclear distribution of DAPI (left), LADs ^(m6^A-Tracer, middle) and transcriptionally active chromatin (H3K27ac, right). **(K)** Mean Euclidian distance distribution from the DINE of H3K27ac : DAPI-(left) and H3K27ac : ^m6^A-Tracer -ratios (right). **(L)** Pearsońs correlation coefficient (PCC) analysis of the distance distribution profiles of H3K9me3, H3K27ac and ^m6^A-Tracer. Graphs in C, E, H, I and K indicate mean ± SD, and n=three independent biological replicates in all experiments. In B, D, G and J pseudocolored Inferno lookup table is used to represent intensity value distributions. Scale bars, 2 µm.

We then examined direct nuclear lamina-chromatin interactions via lamin-associated domains (LADs) (*23*), by tagging LADs in living cells using a DNA adenine methyltransferase (Dam) identification system (DamID). In this system, a lamin-A/C-Dam fusion protein methylates adenine nucleotides in purine ring position 6 (m^6^A) in chromatin adjacent to the nuclear lamina (*65*). This methylated chromatin is visualized using a m^6^A binding protein fused to a red fluorescent protein mScarlet (m^6^A-Tracer-mScarlet) (Fig. 5F). We quantified intensity profiles of LADs (m^6^A-Tracer-mScarlet signal) and repressive (H3K9me3) and active (H3K27ac) histone modifications as a function of distance (0-600 nm) from the DINEs and nucleus edge as described above. The data were similarly normalized to the DAPI signal to account for local chromatin density. We then correlated the m^6^A -Tracer-mScarlet signal with the histone markers to assess the transcriptional status of the LADs. Although the absolute m^6^A-Tracer (LAD-associated) fluorescence intensity at DINEs was higher than at the nuclear edges, the DAPI-normalized intensity was similar in both locations (Fig. 5G and 5H), indicating similar lamina-LAD binding in both locations. The repressive H3K9me3 : DAPI ratio was nearly identical at DINEs (1.17 ± 0.19) and at the edge (1.13 ± 0.53) (Fig. 5G and 5I). In contrast, the active H3K27ac : DAPI ratio was higher at DINEs (0.77 ± 0.21) than at the edge (0.41 ± 0.30) (Fig. 5J and 5K). Interestingly, the association of LADs with H3K9me3 and H3K27ac differed between DINEs and the nuclear edge. At the edge, LAD fluorescence profile correlated strongly with H3K9me3 (Pearsońs correlation coefficient, PCC = 0.93), but not with H3K27ac (PCC = −0.14). However, at DINEs, the LAD fluorescence profile correlated with both H3K9me3 (PCC 0.59) and H3K27ac (PCC 0.77) (Fig. 5L). Taken together, these findings indicate that DINEs are functionally distinct regions of the NE that likely contribute to the spatial and functional 3D organization of chromatin.

### DINE-containing nuclear phenotype correlates with suppression of growth and MAPK-signaling

During epithelial maturation, cells undergo a reversible developmental process that reaches an epithelial state, in which epithelium-to-mesenchymal transition (EMT) is suppressed. This transition is controlled by contact inhibition, which evokes cellular growth arrest and limits proliferation. Contact inhibition in epithelium is established by Wnt/b-catenin, SRF, Hippo, and mitogen-activated protein kinase (MAPK) signaling. In MDCK II cells, the classical MAPKs ERK1/2 (extracellular signal-regulated kinase 1/2) act as the main regulators of contact inhibition (*66–68*, *68*, *69*). Contact inhibition deactivates ERK1/2 via dephosphorylation and transcriptional downregulation of early-response genes. Based on our observation that DINE-formation increases with epithelial maturation and cell density, which both are MAPK/ERK pathway-regulated processes, and decreases during EMT activation in wound healing, we examined this relationship with total mRNA sequencing (mRNA-seq) in newly seeded (2 h) non-DINE-containing cells and mature (7 d) DINE-containing epithelium (n=two independent biological replicates).

The mRNA-seq detected 12 978 commonly expressed genes, 498 unique to the newly seeded, and 280 unique to mature cells (Fig. S4A). Analysis of differentially expressed genes (DEG) revealed 1081 downregulated and 175 upregulated genes in the mature (7 d grown) epithelium (absolute log2 fold change (Abs(log2FC)) >1 and adjusted p-value (p_Adj_) <0.05) (Fig. 6A). To verify that the 7 d sample was epithelial dominant and not mesenchymal, we performed Gene Set Enrichment Analysis (GSEA) using Log2FC values as a pre-ranked list and the Molecular Signatures Database EMT hallmark gene list (*70*). The genes were mostly negatively enriched (FDR q-value = 0.065), indicating EMT suppression in the 7d sample (Fig. S4B). QIAGEN Ingenuity Pathway Analysis (IPA, QIAGEN Inc.(*71*)) showed 187 upregulated and 5 downregulated pathways in the mature epithelium ((-log(p-value)) > 1.3 and |z-score| >2), Table S2). The statistically significant downregulated pathways included “Mitotic Prometaphase”, “RHO GTPase Cycle”, and “Cell Cycle Checkpoints”, consistent with maturation and contact-induced growth arrest of 7 d cells. The IPA analysis identified the Nuclear-Cytoskeleton Signaling Pathway as strongly downregulated ((-log(p-value): 15.10; z-score: −5.56) (Fig. 6 B). This pathway includes cadherins, actomyosin cytoskeleton, NE transmembrane proteins, NPCs, LINC complexes, and nuclear lamins. This indicates that the nuclear mechanotransduction machinery from the plasma membrane to the NE is significantly affected during epithelial maturation.

**Fig. 6.**
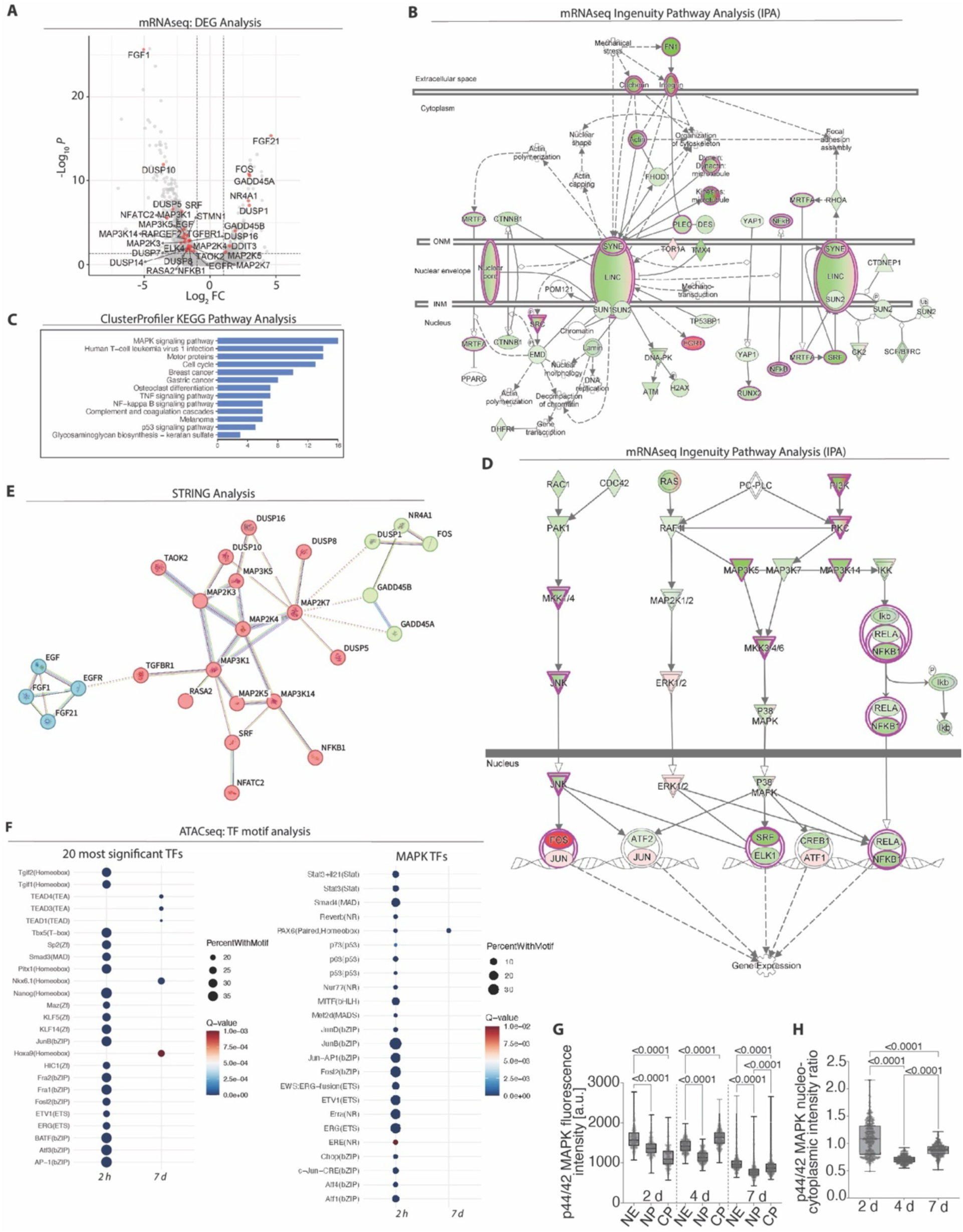
Nuclear mechanosensing and MAPK-signaling are suppressed in DINE-containing mature epithelium. **(A)** The volcano plot of the differentially expressed genes (DEGs) in mature 7 d grown epithelium in comparison to developing 2 d grown epithelium. MAPK-signaling -associated genes highlighted in red. **(B)** Ingenuity pathway analysis (IPA) pathway generator -derived map presenting affected nuclear mechanotransduction and NE -associated components and their downstream actors. Red and green indicate up- and down-regulation, respectively. Purple lines indicate statistically significantly affected components (Abs(log2FC)) >1 and pAdj <0.05). **(C)** ClusterProfiler KEGG pathway over-representation “MAPK signaling pathway” analysis of the 7 d -grown mature epithelium showing significantly affected biological processes in the 7-d grown mature epithelium in comparison to the 2 d-grown epithelium. **(D)** IPA pathway generator -derived map showing affected components of MAPK-signaling. Red indicates up-regulated, and green indicates down-regulated. Purple lines indicate statistically significantly affected components (Abs(log2FC)) >1 and pAdj <0.05). **(E)** STRING analysis of MAPK-associated DEGs in the 7 d grown epithelium, indicating functional enrichment in three clusters within interconnected networks. **(F)** Transcription factor (TF) motif analysis of transposase-accessible chromatin with high-throughput sequencing (ATAC-seq) of 2 d and 7 d grown cells indicates global decreased accessibility of TF motifs, highlighting significant suppression of MAPK/ERK-associated motifs in the mature epithelium. Box-and-whiskers plots presenting **(G)** intracellular normalized fluorescence intensity distribution, and **(H)** nucleo-cytoplasmic ratio of activated phosphorylated ERK1/2 (phospho-ERK, p44/42 MAPK) at 2, 4, and 7 d post-seeding. One-way ANOVA with Dunnett’s multiple comparisons test indicates statistical significance, n=three independent biological replicates. Box-and-whisker plots represent the 25^th^–75^th^ percentiles as boxes, the median as a line within the box, and whiskers indicating the minimum and maximum values.

In differential expression analysis of the 248 genes in the MAPK-pathway (MAPK Signaling Pathway from Harmonizome 3.0, downloaded 28.2.2025)(*72–74*), 181 were found from the dataset with 8 genes significantly upregulated and 24 genes significantly downregulated in the 7 d grown cells (Fig. 6A and Table S3).) The upregulated genes included DUSP1 (*75*), which is an inhibitor of EMT and MAPK (ERK, p38, and JNK) signaling, and growth arrest inducible GADD45A and GADD45B. Downregulated genes included multiple MAPK kinase kinases (MAP3Ks), MAPK kinases (MAP2Ks), and their regulators and effectors (DUSP10, PDGF, DUSP5, EGF, FGF1) (Fig. 6A). ClusterProfiler KEGG pathway over-representation analysis using “MAPK signaling pathway” showed significant enrichment (Benjamini p=1.6E-2, 4.8 % i.e., 32 genes) with 47 significantly upregulated and 25 downregulated MAPK-associated genes (Fig. 6C and Table S3). IPA core analysis confirmed suppression of MAPK-pathways, as significantly downregulated pathways included p38 MAPK (-log(p-value): 3,22; z-score: −2,14); RAF-independent MAPK1/3 activation (-log(p-value): 7,56; z-score: −2,53); and RAS-RAF-MEK-ERK cascade (-log(p-value): 4.06; z-score: −4.71) (Table S3). To generally visualize the most common MAPK routes, we used the IPA pathway generator to map mRNA-seq results (Fig. 6D). We also performed a Search Tool for the Retrieval of Interacting Genes/Proteins (STRING) database analysis (medium confidence threshold score ≥ 0.4, disconnected nodes excluded) (*76*) on the MAPK-associated genes and identified common interaction networks (Fig. 6E and Table S4). Together, these findings suggest contact inhibition-mediated growth arrest and network-wide suppression of the MAPK-mediated pathways in the mature DINE-containing epithelium.

Next, we analyzed chromatin organization during MAPK suppression by mapping accessible chromatin regions in the mature and the newly seeded (2 h) epithelium using ATAC-seq, followed by transcription factor (TF) motif analysis with a focus on downstream MAPK targets. ATAC-seq showed that most chromatin in 7 d DINE-containing nuclei was transcriptionally inaccessible, with few accessible regions mapping mainly to distal intergenic sequences (95.11 %, Fig. S4, C to E). TF motif enrichment analysis indicated significant changes between the differentially accessible peaks in 2 h and 7 d cells, with 13 motifs significantly enriched in peaks differentially accessible in 7 d vs 2 h, while 294 were significantly enriched in peaks differentially accessible at 2 h vs 7 d (Benjamini-Hochberg q<0.01) (Table S5). Focusing on the motifs present in more than 15 % of the peaks, TEAD-family members, Nkx6.1, and Hoxa9 target motifs were overrepresented in the peaks that were more accessible in the 7-d grown sample (Fig. 6F). Less accessible motifs included Fra1/2, Atf3, KLF5, AP-1, and Ets family members (ETV1, ERG), which are common targets of MAPK/ERK, p38 MAPK, and JNK, consistent with overall downregulation of MAPK-signaling (*77–82*). In total, 23 of MAPK-associated TFs were found to be downregulated in 7d cells in comparison to the 2 h cells (Fig. 6F and Table S5). These ATAC-seq results support wide downregulation of the MAPK signaling at the chromatin level in mature DINE-containing cells.

To determine the importance of MAPK/ERK signaling in the contact inhibition and maturation of epithelial cells, we sought to decipher the correlation of MAPK/ERK inhibition and epithelial maturation. Once MAPK/ERK signaling becomes inactive, dephosphorylated ERK1/2 localizes predominantly to the cytoplasm and becomes sequestered at the NE by binding to A-type lamins (*83*). Upon MAPK activation, ERK1/2 is phosphorylated, relocates into the nucleus, and the nuclear lamina-bound ERK1/2 detaches from the lamin A to activate transcription. We next analyzed the subcellular distribution and quantified the fluorescence intensity and nucleo-cytoplasmic intensity ratio of active phosphorylated ERK1/2 with phospho-ERK1/2 (p44/42, Thr202/Tyr204) antibody in cells grown for 2, 4, or 7 d (Fig. 6G and 6H). The phospho-ERK1/2 signal decreased over time, with lower fluorescence intensity at the NE, nucleus, and cytoplasm from 2 d to 7 d, supporting that MAPK signaling was inhibited in the mature DINE-containing cells (Fig. 6G). Specifically, mean intensity of phospho-ERK1/2 in 2 d and 7 d grown cells decreased from 1620 ± 20 a.u. (mean ± SEM) to 1010 ± 20 a.u. at the NE, 1400 ± 20 a.u. to 800 ± 10 a.u. in the nucleoplasm, and 1140 ± 20 a.u. to 920 ± 20 a.u. in the cytoplasm, respectively. The nucleo-cytoplasmic intensity ratio of phospho-ERK1/2 was significantly higher in 2 d grown cells, indicating nuclear enrichment (Fig. 6H). At 4 d, and even more pronounced at 7 d, nuclear phospho-ERK1/2 levels were significantly reduced (Fig. 6G). Altogether, the nucleo-cytoplasmic ratio of phospho-ERK1/2 decreased from 1.10 ± 0.02 (2 d) to 0.7 ± 0.07 (4 d) and finally to 0.90 ± 0.01 (7 d) (Fig. 6H). These results revealed a clear reduction in the ERK1/2 activity in mature DINE-containing cells. Together with our sequencing data, these results further support that MAPK inhibition correlates with the presence of DINEs in the mature epithelium.

### Mechanical lateral compression induces DINE-formation and correlates with MAPK suppression

Our results suggested that downregulation of MAPK signaling is associated with DINE formation in the mature epithelium. We therefore tested whether increased cell packing of near-mature epithelium by lateral compression can directly inhibit MAPK signaling and induce DINE formation. For this, we used mechanical manipulation of cells by lateral compression and quantified the proportion of DINEs and transcriptional EMT and MAPK-related changes with mRNA-seq before and 2 h after the compression.

We established the lateral compression using our in-house -developed Brick Strex cell manipulation device (*84*). Cells stably expressing histone H2B-mCherry or LA-CB-EGFP were grown for 3-4 d to form an intact near-mature monolayer on a pre-strained PDMS membrane sheet pre-stretched by 25 % strain (Fig. 7A). The uniaxial lateral compression was achieved by gradual relaxation of the membrane, reducing the surface area under the cells, and introducing −20 % lateral compressive strain and higher cell density (*84*). After a 2 h recovery, cells were fixed and analyzed for the cell density (nuclei/µm^2^) and proportion of DINE-containing nuclei. Similarly, cell density and DINEs were quantified from non-compressed control cells. In the compression, the cell density increased from ~3290 ± 1180 to 4280 ± 1040 cells/mm^2^, (~30 % increase, Student’s unpaired T-test, p=0.0292) (Fig. 7B). The proportion of DINE-containing nuclei increased from ~28 % (non-compressed control) to 42 % (after compression) (1.5-fold increase, Student’s unpaired T-test, p=0.0003; Fig. 7B). These results show that lateral compression of a near-mature monolayer directly induces DINE formation.

**Fig. 7.**
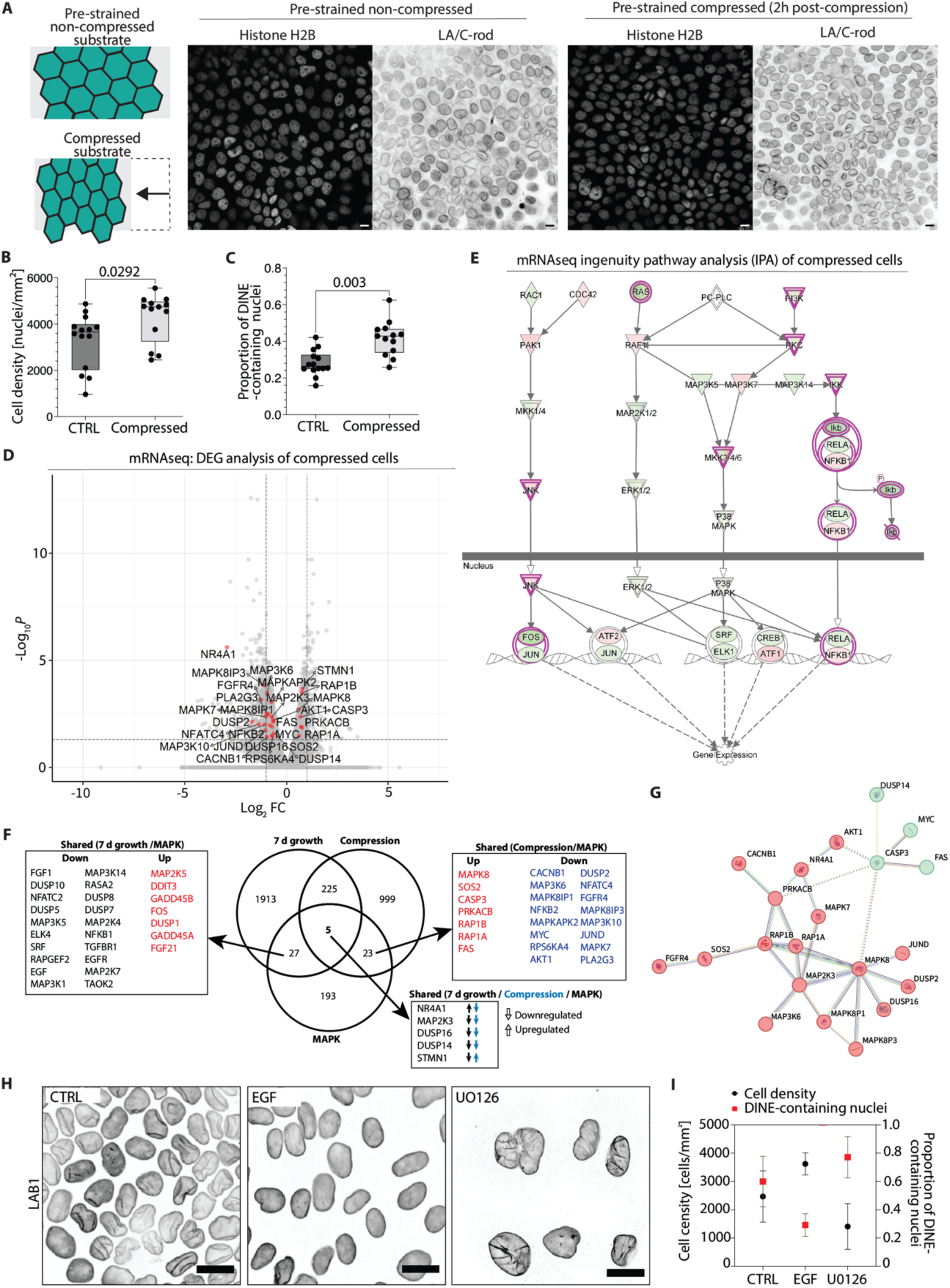
Suppression of MAPK signaling induces DINE-formation. **(A)** Schematic representation of lateral compression experiment showing changes in epithelial monolayer morphology and cell density in non-compressed and compressed states. Box-and-whiskers plots showing quantification of **(B)** cell density and **(C)** proportion of DINE-containing nuclei in the population before and after compression, n=three independent biological replicates. Box-and-whisker plots represent the 25^th^–75^th^ percentiles as boxes, the median as a line within the box, and whiskers indicating the minimum and maximum values. **(D)** Volcano plot of the differentially expressed genes (DEGs) in compressed epithelium recovered for 2 h in comparison to 2-3 d grown non-compressed epithelium. MAPK-signaling-associated genes highlighted in red. **(E)** Ingenuity pathway analysis (IPA) pathway generator -derived map presenting affected MAPK-signaling pathways and down-or up-regulation of their downstream actors. Red and green indicate up- and down-regulation, respectively. **(F)** Venńs diagram indicating uniquely expressed or shared DEGs in 7d grown mature epithelium, compressed epithelium, and their association with MAPK pathway genes. **(G)** Molecular interaction and directionality of regulation STRING cluster analysis of MAPK genes detected in 7 d grown cells. K-means clustering identifying three clusters. **(H)** Lamin B1 (LB1) immunostaining of non-treatedand EMT driver EGF or ERK1/2 inhibitor U0126 -treated monolayers 5 days after treatment. Scale bars, 10 µm. **(I)** Quantification showing the cell density and proportion of DINE-containing nuclei in non-treated control and drug-treated cells grown for 5 d. Data are represented as mean ± SD, n=three independent biological replicates.

To assess the EMT status (maturity) and MAPK-related changes before and 2 h after the compression, we performed mRNA-seq and differential expressed genes (DEG) analysis. We detected 12 585 common genes, with 793 unique to non-compressed and 141 unique to compressed samples (Fig. S4F). The DEG analysis revealed 1580 significantly downregulated and 1262 upregulated genes (pAdj <0.05). Most of these had Abs(log2FC) < 1, which was expected with brief 2 h post-compression recovery. Therefore, in the further analyses, we applied a threshold of Abs(log2FC) ³ 0.585, thus over 1.5-fold change in the expression. Similarly to the 7 d grown cells, Ras-Raf-MEK/ERK signaling was downregulated before compression and thus was not enriched in the DEG analysis of compressed cells. These were expected as the 2-3 d grown monolayers are confluent, but not yet mature. In contrast, the GSEA using a generated pre-ranked list based on the fold change of each gene and EMT hallmark set as a gene set (HALLMARK_EPITHELIAL_MESENCHYMAL_TRANSITION, The Molecular Signatures Database (MSigDB)) (*85*) showed stronger negative enrichment in the compressed cells in comparison to the non-compressed control and the 7-d grown cells (Fig. S4B), indicating suppression of the EMT following lateral compression (Fig. S4G). Next, pre-ranked lists were generated based on the expression level of the genes separately from both samples with MAPK-associated genes as a gene set (MAPK Signaling Pathway from Harmonizome 3.0, downloaded 28.2.2025) (*72*, *73*). The GSEA indicated broad expression (positive and negative enrichment) of MAPK-associated genes before and after compression (Fig. S4H). The DEG analysis revealed differential expression of 28 MAPK-associated genes, with 20 downregulated and only 8 upregulated, indicating enhanced repression of MAPK signaling in response to compression (Fig. 7D). The IPA core analysis did not detect statistically significant changes in MAPK-associated pathways following compression, presumably because the non-compressed control already showed active contact inhibition and low LogFC during the 2 h recovery time. However, when the data were mapped into previously generated visualization of MAPK pathways, it demonstrated significant downregulation of Ras, PI3K, MAP2K3/4/6 upstream ERK1/2, p38 MAPK, and JNK, and increased MAPK suppression following the compression (Fig. 7E, Table S2, and Table S3). Comparison of the compressed sample DEGs to those of the mature monolayer revealed that while significantly upregulated in mature epithelium, the compression did not induce significant upregulation of growth arrest genes or negative regulators of MAPK-signaling (GADD45A, GADD45B, DUSP1) within the observed time scale (Fig. 7F). The comparison showed only 5 overlapping MAPK-related genes between the conditions. These included MAP2K3 (p38 MAPK), DUSP16, DUSP14, NR4A1, and STMN1 (Fig. 7F). Of these, expressions of MAP2K3 and DUSPs had changed in the same direction and were downregulated in both cases.

To explore protein-protein interactions (PPI) and identify functional clusters related to these 5 common genes, we performed STRING analysis (version 12.0, medium confidence threshold score ≥ 0.4, disconnected nodes excluded) without identifying any biologically relevant interactions. The STRING analysis (31 nodes, average node degree 2.45, PPI enrichment p-value < 1.0e-16) from all MAPK-associated DEGs in the 7 d grown mature epithelium (27 unique and 5 commonly expressed genes, Fig. 7F) indicated three distinct clusters with JNK and ERK pathway-activator MAP3K1 as a central affected gene within the MAPK-signaling pathway cluster (Fig. 7G). The analysis also identified several MAP2Ks (MAP2K3, MAP2K4, MAP2K7), which indicated suppression of JNK and p38 MAPK signaling. In contrast, from all MAPK-associated DEGs in the compressed cells (23 unique and 5 commonly expressed, Fig. 7F), the STRING analysis (28 nodes, average node degree 2.07, PPI enrichment p-value 1.11e-16, k-means clustering) revealed two distinct clusters, with MAP2K3 positioned centrally within a module enriched for the MAPK-signaling pathway, suggesting a p38 MAPK-mediated response. Together, these results suggest that while both maturation and mechanical compression induce DINEs, these processes may involve early contact inhibition and distinct regulation of MAPK signaling depending on the type and time scale of the associated mechanical cues. Together, these results imply the importance of MAPK suppression in the DINE-containing cells.

### Suppression of MAPK/ERK signaling induces DINE-formation independently of cell density

Finally, as our results indicated suppression of MAPK/ERK both in the DINE-containing mature epithelium and in the non-mature epithelium before and after lateral compression, and because of its role in contact inhibition governing epithelial maturation in MDCK II cells (*68*), we sought to identify its role in DINE formation by inhibition of MEK1/2 to block ERK1/2. Non-treated control cells were grown for 5 days, or for 2 days followed by treatment with either the ERK1/2 activator epidermal growth factor (EGF, 100 ng/mL) or the MEK1/2 inhibitor U0126 (50 µM) (*86*) for 3 d. After this all cells were fixed, immunostained for LB1 and following LSCM imaging quantified for their proportion of DINE-containing nuclei. First, we assessed if MEK1/2 inhibition alters the localization of phospho-ERK1/2, as shown in experiments in Fig. 4F. The nontreated control cells and drug-treated cells were fixed 3 d post-seeding and immunostained for phospho-ERK1/2 and LA/C-N. LSCM showed nuclei without DINEs in the control and the EGF-treated monolayers, with phospho-ERK1/2 distributed to the nucleus and cytoplasm. In contrast, the nuclei of U0126-treated cells displayed DINE-like invaginations, and phospho-ERK1/2 was enriched in the cytoplasm, at the NE, and markedly, within the DINE-like invaginations, consistent with ERK1/2’s known sequestration within the nuclear lamina during MAPK suppression (*83*) (Fig. S5B). To quantify the influence of MAPK inhibition on nuclear DINE-containing phenotype, cells were similarly treated 24 h post-seeding with either U0126 or EGF (100 ng/mL) to either inhibit or promote growth factor-mediated cell proliferation, respectively, and to allow development of the monolayer. Cells were fixed at 5 d post drug treatment, alongside their non-treated controls, to avoid cytotoxic effects of the drugs. Cells were immunostained for LB1 and DAPI to analyze the presence of DINEs. Quantification of DINEs based on the DAPI and LB1 staining showed that ~60 % of the nuclei in non-treated control cells contained NE deformations (wrinkles) resembling DINEs, i.e. invaginations that visually extended inward and longitudinally by at least one half of the apparent nuclear diameter and were associated with chromatin. EGF activation reduced the proportion of nuclei with DINE-resemblance to ~29 %. In contrast, inhibition of the ERK1/2 with U0126 increased the fraction of these nuclei to ~80 %, despite a significantly lower cell density in comparison to the untreated control cells (Fig. 7, H and I). Together, these results support our previous findings and suggest that while other pathways may also contribute to the process, MAPK/ERK inhibition is sufficient to induce NE invagination. Together, these findings highlight that DINEs are not merely a mechanical response to forces but also reflect cellular identity and signaling state.

## Discussion

Biomechanical guidance plays a critical role in cellular function, aging, and the pathogenesis of various diseases, including muscular dystrophies, cardiomyopathies, fibrosis, and cancer (*56*, *87*, *88*). In cancer, force-mediated changes in the tumor microenvironment and intercellular communication drive cellular plasticity and reprogramming, promoting invasiveness through changes in drug resistance, proliferation, and metastatic potential (*89–94*). Through mechanotransduction, physical cues originating from the external microenvironment or intrinsic cellular processes affect nuclear shape and molecular architecture by deforming the NE (*95*). At the NE, nuclear lamina and chromatin define the intrinsic mechanical properties of the nucleus, and interconnect chromatin with the NE, allowing mechanogenomic effects. Force-driven NE deformation is also accompanied by biochemical signaling that co-regulates transcription of genes via mechanosensitive transcription factors, underscoring the significant role of NE deformation in cell physiology. Complex nuclear morphologies have been previously acknowledged and documented (*96*). Among these features, the nucleoplasmic reticulum (NR) has been described as invaginations traversing through the nucleus linked to SUN1 and cytoskeletal filaments (*31*, *43*). From another hand, NE invaginations have been considered as potential artefacts of sample preparation and because of this, reporting of these nuclear phenotypes and their broader recognition have presumably been limited. Nevertheless, invaginated nuclei have already been documented in the electron microscopy images from 1970’s histology books and contemporary resources like Cell Atlas OpenOrganelle (*97*, *98*), suggesting that these structures are genuine and undercharacterized. Recent models propose that the NE can adopt diverse morphologies with minimal mechanical resistance, particularly in crowded environments such as tumor microenvironments, where these shapes may correlate with cancer aggressiveness and invasiveness (*38*, *41*). A deeper understanding of the NE invaginations is therefore essential for refining interpretations of nuclear morphology in both health and disease.

Despite advances in molecular diagnostics, nuclear morphology remains a cornerstone in histopathology and oncology, where abnormal nuclear shapes, reduced LA/C expression, and altered lamin-mediated nuclear mechanics are considered to associate with malignancy and aging (*38*, *63*, *99*, *100*). Because over 90 % of cancers are carcinomas, i.e., of epithelial origin, we sought to deepen our understanding of epithelial nuclear morphology during the growth and maturation of a healthy epithelium. We found that nuclear morphology can be significantly diverse in the epithelium also in non-pathological contexts. We present evidence of deep invaginations of the nuclear envelope (DINEs) in healthy mature epithelium. This challenges conventional diagnostic paradigms, and broadens our understanding of nucleus shape control, the role of NE in spatial chromatin organization and cellular transcriptional identity, and the potential importance of nuclear lamina in mechanogenomics.

Our study reveals DINEs as prevalent components of epithelial nuclear architecture *in vitro* and *in vivo*, potentially contributing to the spatial and functional organization of the genome during epithelial maturation or in response to mechanical stimuli such as cellular crowding. We observed DINEs in epithelial cells *in vivo* in mouse esophagus and duodenum, as well as in human esophageal organoids, indicating that these structures are intrinsic to epithelial cells and not artifacts of *in vitro* culture or sample processing. Using LSCM and high-resolution imaging techniques ExM and TEM, we found that over 90 % of the nuclei in confluent mature epithelial monolayers *in vitro* exhibited DINEs. These structures were enriched in densely packed chromatin, suggesting a functional role in spatial chromatin organization.

The DINEs existed independently of actomyosin cytoskeletal tension or nuclear connections to the cytoskeleton, implying that DINEs may be maintained by an intranuclear mechanism, potentially involving chromatin interactions. Our investigation into the role of A-type lamins revealed their necessity for DINE formation and normal nuclear morphology. *LMNA* deletion resulted in aberrant nuclear shapes and loss of DINEs, while overexpression of LA/C altered nuclear architecture and DINE prevalence, indicating that A-type lamins are critical for maintaining these structures. Quantification of lamins showed that the DINEs contained basal NE -like nuclear lamina organization with significantly increased LA/C epitope accessibility. The DINE formation was closely associated with increased epithelial cell density, EMT, contact inhibition-associated growth arrest, and maturation. Time-course experiments demonstrated that DINEs emerge following cellular crowding, likely in response to increased cellular packing, contact inhibition, and epithelial maturation. Observations during wound healing further supported this, showing that DINE-formation correlated well with local density variations, and EMT suppression. Moreover, we demonstrated that DINEs dynamically remodel during confined 3D migration, potentially facilitating epithelial nuclear deformation in constricted environments. Previous studies have suggested that excess nuclear lamina surface area facilitates migration through tight spaces (*101*); our findings extend this by showing that DINEs may unfold to accommodate such deformation, enabling cells to adapt to mechanical constraints within a tissue, for example, during wound healing.

We also explored the functional implications of DINEs in chromatin organization. Prior research has shown that chromatin status and its NE interactions can influence nuclear morphology, whereas changes in nuclear volume affect the electrostatic potential and surface area to volume ratio of the nucleus, thereby modulating nuclear lamina-associated chromatin and epigenetics in the epithelium (*102–104*). Our data revealed that DINEs are associated with chromatin regions of active transcription, marked by phosphorylated RNA polymerase II and increased LA/C epitope accessibility. This suggests a role for DINEs in organizing chromatin and transcriptional activity. Furthermore, our ATAC-seq analysis indicated increased accessibility of intergenic regions, implying that DINEs may differentially facilitate the transcriptional activation of LADs, a notion supported by H3K27ac enrichment within the DINEs (*105*).

Our total mRNA sequencing further revealed that DINE formation correlates with suppression of MAPK-signaling. In mature epithelium, control of cell proliferation is essential for tissue homeostasis, morphogenesis, wound healing, and often dysregulated in cancer. Epithelial cells regulate this through contact inhibition, which in *in vitro* conditions is affected by E-cadherin - associated adherens junctions interacting with the cytoskeleton and catenins mediating downstream signaling, including Wnt, Hippo-pathway, and the MAPK pathway (*66*, *106*). Consistent with our previous findings that lateral compression redistributes b-catenin to the cell vertices (*84*), we observed that this mechanical modulation of the epithelium leading to DINE formation also significantly suppressed MAPK signaling. This included downregulation of MAPK pathway genes, upregulation of MAPK inhibitors, and reduced ERK1/2 phosphorylation with increased cytoplasmic retention in DINE-containing mature epithelial cells. This aligns with recent studies showing that compressive forces can activate epigenetic programs via MAPK/ERK signaling (*107*).

Additionally, we found that NE invaginations could be induced by MAPK inhibition even at low cell density, whereas EMT activation by growth-promoting factors counteracted DINE formation. Our data also highlight that increased mechanical stress through lateral compression promotes both DINE formation and MAPK suppression, particularly via the p38 MAPK pathway. These findings suggest that epithelial maturation and mechanical compression may suppress MAPK through distinct mechanisms, underscoring the role of negative MAPK regulation in the mature epithelium. Our results propose that DINE formation may be part of a broader regulatory network downstream of MAPK signaling, contributing to the spatial and functional organization of chromatin downstream of mechanical stimuli.

In conclusion, our study provides new insights into the structural and functional significance of DINEs in epithelial cells. We demonstrate that DINEs exhibit a basal-like nuclear lamin organization with distinct LA/C accessibility, and unique chromatin features, with their formation correlating with MAPK suppression. By integrating extracellular mechanical cues, DINEs may modulate gene expression and coordinate cellular responses, highlighting the importance of nuclear architecture in epithelial phenotype regulation. To the best of our knowledge, this is the first study where the dynamics and functional role of DINEs, their interconnected nuclear lamin and chromatin organization and their implications on transcriptional profiling have been quantified. This work provides conceptual advancement in our understanding of nuclear membrane deformation by highlighting DINEs as nuclear regulatory elements with functional consequences for cellular behavior and phenotype conversion. Collectively, our findings open new avenues for investigating the mechanisms that govern epithelial homeostasis through nuclear morphology and lamin-mediated mechanogenomics in the ever-changing cellular microenvironment in our tissues.

## Materials and Methods

### Experimental Design

The aim of this study was to delineate the mechanistic basis underlying the formation and persistence of deep invaginations of the nuclear envelope (DINEs) in epithelial cell nuclei, and to elucidate their functional significance in modulating intranuclear chromatin organization and function, and in co-regulating cellular phenotypic behavior. The experiments were done *in vitro* by using a traditional epithelial mechanobiology model cell line, state-of-the-art fluorescence and super-resolution confocal imaging, 2D and 3D mechanical manipulation of cells, micropatterning, 3D microfluidics, mouse and human 3D organoids, total mRNA and ATAC-seq sequencing, and imaging methodologies ATAC-see and DamID enabling combining fluorescence imaging with sequencing. Presence of invaginated nuclei was also validated *in vivo* using mouse tissue duodenum and esophagus tissue sections.

### Cells

Madin-Darby canine kidney (MDCK) type II and its derivative LA/C KO cell line (*58*) were maintained in low glucose MEM (#41090093, Thermo Fisher Scientific, Gibco^TM^, Paisley, UK) supplemented with 1 % (vol/vol) penicillin-streptomycin antibiotics (P/S, #15140122, Thermo Fisher Scientific, MA, USA) and 10 % fetal bovine serum (FBS, #10500064, Thermo Fisher Scientific). MDCK II cells expressing lamin A/C-Dam methylase (LA/C-Dam) and m6A-Tracer-mScarlet were maintained in complete medium without P/S antibiotics and with Geneticin (0.25 mg/mL, G418 sulfate, #10131035, Gibco, Thermo Fisher Scientific) and Zeosin (100 µg/mL, R25001, Gibco, Thermo Fisher Scientific). LA-EGFP cells were maintained in MEM with P/S and Geneticin (0.25 mg/mL). Cells were maintained at standard 37 °C in a humidified atmosphere with 5 % CO_2_ and passaged once a week. For the experiments, cells were seeded (1.2 × 10^6^ cells/mL, 100 µL) on collagen I (#A1064401, Thermo Fisher Scientific) -coated high-performance cover glasses (D = 0.17 mm, Carl Zeiss Microscopy, NY, USA) of size 18×18 mm^2^ for immunofluorescence LSCM and ATAC-see, and 22×22 mm^2^ for ExM and micropatterning.

### Pharmaceuticals

Pharmaceuticals used in the experiments were DMSO (0.2 %, D2650-100ML, Merk/MilliporeSigma, MA, USA), acrylamide (5 mM, 30 % Acrylamide/Bis, #1610154, Bio-Rad, CA, USA), nocodazole (5 uM, FN12330, Biosynth Carbosynth, Compton, UK), EGF (#72528S, Cell Signaling Technology), U0126 (9903S, Cell Signaling Technology), cytochalasin D (5 ug/ml) and Y27632 (10 uM, Y0503-1MG, Sigma Aldrich, Merck).

### Antibodies

A-type lamins lamin A/C were detected with a mouse monoclonal antibody (mMAb) against lamin A/C C-terminus residues aa 319-566 (1:400, LA/C-C, [131C3], ab1791, Abcam, Cambridge, UK) or a rabbit monoclonal Ab (rMAb) against full-length lamin A/C (1:500, LA/C-rod, [EP4520-16], ab133256, Abcam). Lamin B1 was detected with a rabbit polyclonal anti-lamin B1 Ab (1:500, ab16048, Abcam). F-actin was visualized with phalloidin ATTO 633 (1:100, ATTO-TEC GmbH, Germany). Histone H3 and its post-translational tail modifications were targeted with anti-H3 (1:500, ab1791, Abcam), anti-H3K9me3 (1:500, ab8898, Abcam), and anti-H3K27ac (1:500, ab4729, Abcam). Active MAPK was detected by using a mouse monoclonal Ab identifying the phosphorylated p44/42 MAPK (1:400, ERK1/2, Thr202/Tyr204 [E10], #9106, Cell Signaling Technology, MA, USA). To detect transcriptionally active RNA polymerase II, anti-RNA polymerase II CTD repeat YSPTSPS (phospho S2) rabbit polyclonal Ab (1:500, ab5095, Abcam) was used. Nuclear pore complexes were detected with a mMAb Mab414 (1:1000, ab24609, Abcam). Secondary antibodies goat anti-mouse and anti-rabbit IgG conjugated with Alexa 488, 568 or 647 (1:200 in 3 % BSA/PBS, Thermo Fisher Scientific) were used in the detection of the primary Abs.

### Immunostaining

Cells were fixed with 4 % paraformaldehyde (PFA, #15713-S, Electron Microscopy Sciences, Hatfield, PA, USA) for 10 min at room temperature (RT), rinsed twice with 1X PBS, and permeabilized for 10 min in RT using 0.5 % Triton X-100 in PBS supplemented with 0.5 % BSA (bovine serum albumin). Samples were incubated with primary antibodies (Abs) diluted in PBS containing 3 % BSA for continuous blocking (1h at RT) using recommended immunofluorescence labeling working concentrations. Incubation was followed by washing once with permeabilization buffer A, once with PBS, and again with permeabilization buffer A. For detection, the samples were treated with secondary antibodies for 1 h at RT in dark, followed by one PBS wash and one deionized water wash (á 10 min, RT, in dark). Samples were mounted with Prolong Diamond with DAPI (P36965, Thermo Fisher Scientific) and cured o/n in the dark at RT. Confined migration samples were mounted by using Vectashield (Vector Laboratories, CA, USA).

### Light microscopy

In LSCM, unique settings were applied to each sample, followed by imaging with constant laser powers and detection voltages, enabling comparable quantitative analysis of the fluorescence intensities. Microscopy imaging was done with a Nikon A1R+ laser scanning confocal microscope (NIS Elements, v. 5.11) mounted in Nikon Eclipse Ti2-E (Nikon Instruments, Tokyo, Japan) and with Nikon 60X/1.40 Apo λS DIC N2 oil immersion objective with 0.14 mm working distance. Solid-state lasers with excitation wavelengths of 488 nm, 561 nm, and 640 nm were used in excitation. Fluorescence emissions were collected with 525/50, 540/30, and 595/50 bandpass filters, respectively, with laser intensities adjusted to avoid photobleaching. Detector sensitivity was adjusted to optimize the image brightness and to avoid saturation. Laser powers and detector voltages were determined individually per treated antibody pair, and after the initial setting, kept constant for each sample to allow ratiometric imaging and quantitative comparison of the fluorescence intensities within the drug-treated and non-treated control samples. The images were 1024 × 1024 pixels, and the pixel size was 41-52nm, or 104 nm in x/y to achieve Nyqvist resolution. The images were acquired without averaging and by first focusing on the bottom surface of the sample, where the position of the sample stage was set as z0 = 0. The fluorescence signal intensities from all emission channels were then collected from bottom to top as optical z-section series with a 150 nm step size. The analysis was performed in ImageJ FIJI distribution (*108*) by generating maximum intensity projections from the acquired z-section series. In wound healing, images of 2151 × 512 with a pixel size of 0.28 µm in x/y were acquired with 30 min intervals.

### Transmission electron microscopy

MDCK II cells were seeded for 7 days on collagen I-coated 18 x 18 mm coverslips and fixed in 3.75 % paraformaldehyde and 0.25 % glutaraldehyde in 50 mM phosphate buffer (pH 6.8). Following post-fixation in 1 % osmium tetroxide for 1 h on ice, the cells were dehydrated in a graded ethanol series and embedded in low-viscosity resin (TAAB Laboratories Equipment Ltd, UK). Thin sections perpendicular to the cell surface were cut with Ultracut UC6a ultramicrotome (Leica Mikrosysteme GmbH, Germany), collected on Pioloform-coated single-slot copper grids, and stained with 2% aqueous uranyl acetate and lead citrate. A transmission electron microscope JEOL JEM-1400 (JEOL Ltd., Tokyo, Japan) was operated at 80 kV, and it was equipped with a bottom-mounted Quemesa CCD camera of 4008 × 2664 -pixel resolution (EMSIS GmbH, Münster, Germany), which was used to obtain the micrographs.

### Expansion microscopy

Sample expansion was performed using a previously described proExM protocol(*109*). The samples were first fixed and immunostained using standard procedures, and anchored in 1 % 6-([acryloyl] amino) hexanoic acid succinimidyl ester (#A20770; Acryloyl X-SE; Thermo Fisher Scientific) in PBS for over 6 h at RT. After anchoring, the samples were washed twice with PBS (á 15 min). Before the gelation, glass coverslips (18 mm × 18 mm, Zeiss high performance, Carl Zeiss Microscopy) were prepared as a base for gel casting. Parafilm (P7793, Merck) was glued on top of a coverslip with cyanoacrylate glue (Loctite Super Glue Power Flex Gel Control; Henkel Norden AB, Bromma, Sweden), and the parafilm-coated coverslip was moved to a six-well plate lid with parafilm side up, and the parafilm cover paper was removed. An in-house -printed spacer was placed on top of the parafilm to confine the gel to a known thin dimension. The gelation solution was prepared on ice by adding inhibitor (4-Hydroxytempo) and accelerator tetramethylethylenediamine to the monomer solution in 1.5-ml centrifuge tubes (S1615-5500; Starlab International, Hamburg, Germany) with initiator ammonium persulfate (10 % w/v) added just before pipetting the solution. The ready-made gelling solution was vortexed and administered in the middle opening of the spacer. A sample with cells grown as a monolayer was carefully placed on top of the gelling solution, with cells facing the gel, a metal nut was placed on top to ensure thin gels, and the sample was left to polymerize (30–45 min at RT). After polymerization, the parafilm-coated coverslip was carefully removed, and excess gel was trimmed with a scalpel. The gel with the coverslip was transferred to a clean 6-well plate (#150239; Thermo Fisher Scientific). The polymerized gel, together with the sample, was next immersed in a 10-fold volume of digestion solution with 8 U/ml Proteinase K (P8107S; New England Biolabs, Ipswich, MA) in digestion buffer containing 50-mM Tris pH 8.0, 1-mM EDTA (E5134, Merck), 0.5% Triton X-100 (T8787, Merck), and 800-mM guanidine HCl (G3272, Merck) in distilled water (dH_2_O), where it was incubated o/n at RT protected from the light. During incubation, the gel detached and was transferred to a 6-cm dish. The gel was washed 5 times with an excess volume of dH_2_O for 1 h. The expanded gel sample was measured to determine the expansion factor (final dimensions/original dimensions from the spacer opening). A piece of gel was cut with an in-house 3D printed puncher and mounted in a live cell imaging chamber (Aireka Cell, Hong Kong, China) with low-melt agarose (A7431, Merck). The chamber was filled with dH_2_O and imaged immediately with LSCM following 15 min temperature stabilization on the microscopy stage.

### Micropatterning

Micropatterned coverslips with ECM coating were prepared with photoresist lift-off(*110*). Hourglass patterns were constructed on glass coverslips by photolithography, in which UV light is used with a photomask to transfer patterns on a photosensitive coating (photoresist). First, the photomask design was done using AutoCAD® (Autodesk, 2019) by drawing hourglass patterns in a 5 mm x 5 mm area. The photomask was fabricated in a cleanroom facility (Class ISO 6, UV-protected). A cs hardmask blanc (Clean Surface Technology CO., Japan, CBL5009Du-AZ1500) was manufactured using μPG501 Conversion Job Manager (Version 1.8.3, Heidelberg instruments) with exposure time of 22 ms, DEFOC = −2. The mask was developed in a 1:4 mixture of AZ351B/dH_2_O (AZ Electronic Materials, Germany, 10054724960) for 1 min, washed with dH_2_O, and dried under pressurized nitrogen. Etching was done in CHROME ETCH 18 (OSC OrganoSpezialChemie Gmbh, 79600316) for 1 min, and the mask was washed with dH_2_O and dried with pressurized nitrogen. Finally, the mask was exposed to UV light (intensity 25 mW/cm2) in UV-F 400 Vitralit light box (Panacol-Elosol Gmbh) for 1 min, developed again in the previously used AZ351B/dH_2_O solution, washed with dH_2_O, and dried underpressurized nitrogen. To construct the hourglass patterns, glass coverslips (22 x 22 mm, No. 1.5H, Marienfeld, Germany) were cleaned in 2% HellmaneX III (Hellma, Germany) solution in dH_2_O in beakers placed inside a sonicator (model 3800, Branson Ultrasonics) for 30 min, washed with excess dH_2_O and 99% EtOH, and dried with pressurized nitrogen. In the cleanroom, the glasses were rinsed first with acetone, then isopropanol, and finally dH_2_O. The glasses were dried with pressurized air and treated with O_2_ plasma for 2 min using reactive ion etching RIE (Advanced vacuum vision 320 RIE). Then, the positive photoresist S1818 (Microchem) was coated on the glasses (approximately 2 µm layer) in a spin coater (Laurell, model WS-650HZB-23NPPB). Spin-coating was done in three steps: First, 5 s in 2000 rpm (acceleration 520) and then 1 min 30 s in 3800 rpm (acceleration 650). After the spin-coating, the samples were baked on top of a hotplate at 100℃ for 2 min (Torrey Pines scientific Inc., models HS60 and HS61) and allowed to cool down. Exposure was done with an OAI mask aligner (Optical Associates Inc., model J500/VIS, intensity 12 MW/cm2 at 436 nm wavelength, i-line filter installed). Samples were placed in contact with the photomask and exposed for 14 s. After exposure, the samples were developed in MF-319 developer (Microchem) for 1 min, rinsed with isopropanol and dH_2_O, and dried with pressurized air. The patterns were observed with a microscope, and a stylus profiler (DektakXT, Bruker) was used to measure the height of the photoresist layer.

The ECM coating was done following the protocol by Moeller *et al.* (*110*). The S1818 hourglass patterned coverglasses were activated with O_2_ plasma for 15 s at 0.4 mbar, 50 W (Diener Pico plasmacleaner). 0.1 mg/ml of PLL(20)-g[3.5]-PEG(2) solution with 4 % TRITC label (SuSoS, Switzerland) was incubated on the plasma-activated samples for 30 min. The samples were washed with dH_2_O and kept in 1xPBS for hydration. Removal of the S1818 photoresist was done with lift-off. Beakers were filled with different ratios of N-Methyl-2-pyrrolidone (NMP, Sigma-Aldrich) and dH_2_O. After the lift-off, the samples were washed in dH_2_O for 15 min. The lift-off removed the S1818 photoresist from the samples, thus creating glass coverslips with hourglass-s empty, uncoated regions surrounded by PLL-g-PEG-coated non-adhesive regions. The glasses were then coated with 0.1 mg/ml collagen I in 0.02 N acetic acid for 1 h, resulting in ECM-coated hourglass-patterned coverslips suitable for cell culture. MDCKII-Occludin-Emerald cells were passaged on the glass coverslips, and after 1 h, the samples were washed with media to avoid cell adhesion outside the patterns, and let grow for 3-7 days prior to fixing, immunostaining, and microscopy imaging.

### Wound healing

Glass coverslips (0.17 mm, cat. No. 474030-9000-000, Carl Zeiss Microscopy, LLC, Thornwood, NY, USA) were coated with collagen I (0.1 mg/ml in 0.02 N acetic acid) for 1 h at RT, rinsed once with sterile PBS, and placed into 6-well plates. Sterile PDMS stencils were positioned firmly on top of the coated coverslips to create a cell-free gap. MDCK II-EGFP-CB cells were seeded on both sides and grown for 3-7 d until the desired confluency. PDMS stencils were carefully removed using sterile tweezers to generate a well-defined wound between the two opposing monolayers. Wells were gently washed once with pre-warmed culture medium to remove detached cells, and fresh medium was added. Following stencil liftoff, cells collectively migrate into the wound. Wound closure was monitored by live-cell confocal microscopy time-lapse imaging with 30 min intervals. Wound healing was quantified by counting DINE-containing and smooth nuclei over time per field of view across the wound area using ImageJ/Fiji.

### Mechanical manipulation by lateral compression

Lateral compression was performed using our in-house -developed Brick Strex device as described previously(*84*). Here, 2 × 10^4^ MDCK II cells expressing H2B-mCherry or LA-EGFP-CB grown for 2-3 days in columns prepared by cutting the column off from a PET insert (small S device) and attached on a 25 % pre-strained fibronectin-coated (10 µg/mL) thin (0.01“) USP class VI PDMS membrane (8 × 4.5 cm, Specialty Manufacturing, Inc., Saginaw, MI, USA) using high-vacuum silicone grease (Dow Corning, Merck, Darmstadt, Germany). As a control, cells were also seeded on unstretched PDMS membranes in similar columns. Once desired confluency was reached, compression was introduced by gradual relaxation of the PDMS membrane, resulting in a 20 % lateral compression, and cells were allowed to recover for 2 h. After the recovery, cells were fixed in 4 % PFA for 10 min in the dark at RT, washed twice with PBS before immunostaining. After immunostaining, the membranes were cut, mounted on an objective glass, and imaged with LSCM.

### Confined 3D migration in microfluidic devices

Confined 3D migration experiments were performed as described previously (*59*). The microfluidic devices were fabricated using standard soft lithography techniques. SU-8 photoresist molds were patterned to define channel geometries with constrictions ranging from 3–10 µm in width and 5 µm in height. Polydimethylsiloxane (PDMS; Sylgard 184, Dow Corning) was cast onto the molds, cured at 70 °C for 2 hours, and bonded to glass coverslips following oxygen plasma treatment. Devices were rinsed once with Abs. EtOH, once with 1× PBS. Devices were pretreated o/n with fibronectin in 4 °C to allow cellular adhesion, followed by a rinse with 1× PBS and loading of device with medium and stabilization at 37 °C in humidified cell incubator for 30 min. For experiments, 50 000 cells were seeded into the device reservoir and cultured overnight at 37 °C in incubator to allow attachment and migration into the devices. Once the cells had migrated into the entry area in front of the confinements, migration into the devices was initiated by establishing a biochemical growth factor gradient with serum. Gradient was established by pipetting 100 µL of 2 % FBS in medium in the other reservoir with simultaneous pipetting of 100 µL of 10 % FBS in medium to the other. Migration within the constrictions was observed for 24 hours with live cell imaging using Keyence BZ-X810 microscope equipped with a humidified incubator warmed up to 37 °C together with the microscopy stage and objective at 20× magnification using 10-min time intervals (6 fph). Alternatively, devices were prepared similarly for fixation after 24 h migration. Images were acquired with an image size of 960 × 720 pixels with a x/y pixel size of 264.16 µm (0.0104 inch).

### ATAC-see

MDCK II cells (1.2 x 10^6^ cells/mL, 100 µL) were seeded for 2 h or 7 d on collagen I-coated 18 x 18 mm coverslips (0.17 mm, cat. No. 474030-9000-000, Carl Zeiss Microscopy, LLC, Thornwood, NY, USA). Hyperactive Tn5 was produced in-house according to the protocol described previously (see additional “Tn5 transposase expression and purification)(*111*). The Atto-565N -labeled oligonucleotides (oligos, M size, 0.2 µmol) for Tn5 transposome adaptor (Tn5MErev) were designed and purchased from Molecular Probes. The oligo sequences were Tn5MErev, 5′- [phos]CTGTCTCTTATACACATCT-3′; Tn5ME-A-ATTO488, 5′- /5ATTO488/TCGTCGGCAGCGTCAGATGTGTATAAGAGACAG-3′; Tn5ME-B-ATTO488: 5′-/ATTO488/GTCTCGTGGGCTCGGAGATGTGTATAAGAGACAG-3′. The assembly of the Tn5 transposome was performed as described previously(*111*), but we used 50 % of the indicated Tn5 concentration to avoid excessive background. The oligos (Tn5ME-A-Atto488, Tn5ME-B-Atto488, and Tn5MErev) were resuspended in water to a final concentration of 100 µM. Equimolar amounts of Tn5MErev+Tn5ME-A-ATTO488 and Tn5MErev+Tn5ME-B-ATTO488 were mixed in separate PCR tubes (200 µL). The oligo mixtures were denatured in a thermocycler for 5 min at 95 °C and cooled down slowly on the thermocycler by turning off the thermocycler (30-45 min). To perform the transposase labeling reaction, consequently with the latter step a final volume of 100 µL of the transposase labeling solution (SL-Tn5) was prepared (50 µl 2X TD buffer, 1.6 µL of Tn5-ATTO-488N with a final concentration of 100 nM, and adding dH_2_O (48.4 µL) up to 100 µl to produce a SL-Tn5 solution with a Tn5 concentration of 50 µM). This mixture was incubated for 30 min at 37 °C on a preheated heat block. The Tn5 transposome was assembled with the following components: 0.25 vol Tn5MErev/Tn5ME-A-ATTO488 + Tn5MErev/Tn5ME-B-ATTO488 (final concentration of each double-stranded oligo was 50 µM each), 0.4 vol glycerol (100% solution), 0.12 vol 2× dialysis buffer (100 mM HEPES–KOH at pH 7.2, 0.2 M NaCl, 0.2 mM EDTA, 2 mM DTT, 0.2% Triton X-100 and 20% glycerol), 0.1 vol SL–Tn5 (50 µM) and 0.13 vol water. After mixing gently and thoroughly, the solution was left on the bench at RT for 1 h to allow annealing of oligos to Tn5. The cells were fixed with 4 % PFA (cat. No. 15713-S, Electron Microscopy Sciences, Hatfield, PA, USA) for 10 min at RT, rinsed twice with 1X PBS, and permeabilized with lysis buffer (10 mM Tris–Cl pH 7.4, 10 mM NaCl, 3 mM MgCl2, 0.01% Igepal CA-630). After permeabilization, the cells were rinsed twice in 1X PBS and incubated in a humidified chamber box at 37 °C until tranposase labeling reaction. After the transposase reaction, the samples were washed with 1X PBS containing 0.01% SDS and 50 mM EDTA for 15 min three times at 55 °C. After washing, the samples were washed with dH2O three times for 5 min followed by treatment with another permeabilization buffer meant for immunostaining (0.1% Triton X-100, 0.5 % BSA in 1X PBS). Immunostaining was done using standard procedures, and the samples were mounted with Prolong Diamond Antifade with DAPI (Thermo Fischer Scientific). The samples were left to cure at RT in dark o/n and stored at 4 °C prior to microscopy.

### ATAC-seq library generation and sequencing

We collected 50,000 cells and centrifuged them at 500 x g for 5 min, 4°C. Cells were washed once with 50 µL of cold 1xPBS buffer and centrifuged again. Supernatant was removed, and cells resuspended in 50 µL of cold lysis buffer followed by centrifugation with the same settings. Supernatant was removed. ATAC-seq libraries were generated as described earlier (*112*). Briefly, a transposition mix (25 μL 2× TD buffer, 2.5 μL transposase (Tn5, 100 nM final), 22.5 μL water) was added to the nuclear pellet. Reaction was incubated at 37 °C for 45 minutes and amplified using PCR. Samples were purified using Qiagen MinElute PCR Purification Kit and again using Agencourt AMPure XP magnetic beads. Samples were sequenced using Illumina NextSeq high output 2×75 bp settings.

### ATAC-seq data processing

The quality control was done with FastQC (v.0.11.9)(*113*). After trimming the adapters and low-quality reads with Trim Galore (v.0.6.7)(*114*, *115*), reads were aligned to the CanFam3.1 reference genome with Bowtie2 (v.2.4.8)(*116*). Duplicate reads were removed with Picard MarkDuplicates (v.2.27.1), after which the peak calling was conducted using MACS2 (v.2.2.7.1) with the following parameters: macs2 callpeak -g --nomodel --nolambda --extsize 200 --shift -100 -f BED -q 0.05 --cutoff-analysis --call-summits --keep-dup all. The consensus peak set was computed using an iterative removal process described in Corces et al., 2018(*117*), from peak summits extended to a fixed width of 501 bp, ignoring peaks extending beyond the chromosome ends, those with a score per million value < 5, and those observed in only one replicate. Quality controlwas performed using the ataqv toolkit. Differential accessibility testing was conducted in R (v4.3.0) using csaw (v.1.36.1)(*118*) as described in Reske et al., 2020(*119*), applying Trimmed Mean of M-values (TMM) normalization. ChIPseeker (v. 1.38.0)(*120*) was used to assign peaks to genomic features, and Hypergeometric Optimization of Motif Enrichment (HOMER, v. 4.10)(*121*) was used for motif enrichment analysis.

### Dam-ID with m^6^A-Tracer-mScarlet detection

MDCK II cells expressing inducible lamin A/C-Dam methylase (LA/C-Dam) were induced with doxycycline (1 µg/mL, D9891, Sigma-Aldrich, Merck) and incubated for 24 h prior to fixing with 4 % PFA (10 min, RT, in dark). After fixation, cells were rinsed twice with 1xPBS and proceeded to immunostaining.

### Total mRNA-seq

Total mRNA was extracted from MDCK II wt cells using RNeasy Midi Kit (#75144, Qiagen, Hilden, Germany) according to the manufacturer’s instructions. RNA quantity and integrity were assessed using the 5200 Fragment Analyzer (Agilent Technologies, CA, USA). Samples were sent to Novogene Bioinformatics Technology Co., Ltd. for sequencing. In brief, the mRNA was purified by using poly-T oligo -attached magnetic beads. After fragmentation, the first strand cDNA was synthesized using random hexamer primers, and the second strand using the dUTP method. The libraries were checked with Qubit and real-time polymerase chain reaction. Quantified libraries were pooled and sequenced on the Illumina platform using paired-end 150 bp reads (Illumina Sequencing PE150 Illumina), generating approximately 6 Gb of raw data per sample. Base calling was done using Illumina software (CASAVA base calling), generating raw FASTQ files. An index of the reference genome (domestic dog CanFam_3.1) was built, and gene-level counts were generated using Hisat2 v2.0.5 and paired-end clean reads were aligned to the reference genome. For bioinformatics analysis, raw data were processed through Novogene’s in-house Perl scripts, and the data were quality checked with FastQC. Relative expression levels were normalized based on gene length and sequencing depth (fragments per kilobase of transcript per million mapped reads, FPKM). Differential gene expression (DEG) analysis was performed with DESeq2(*122*). Downstream functional enrichment analyses were conducted using GO (PANTHER/GO Ontology database (DOI: 10.5281/zenodo.6399963 Released 2022-03-22), KEGG (Database for Annotation, Visualization, and Integrated Discovery (DAVID), https://davidbioinformatics.nih.gov/), and Gene Set Enrichment Analysis (GSEA)(*123*, *124*) to identify significantly affected biological pathways. GSEA was performed using GSEA software version 4.3.3(*124*, *125*). Preranked lists were generated based on gene expression level or log2FC values, and the analysis was run with the GSEAPreranked tool (1000 permutations, weighted enrichment statistics). VENN diagrams were generated at first using interactiVenn tool(*126*) and modified using Adobe Illustrator. The IPA Core analysis was performed, and the pathways were generated with QIAGEN IPA Interpret (QIAGEN Inc., (*127*, *128*). Canonical pathways, upstream regulators, and molecular networks were identified based on IPA’s curated knowledge base. Volcano plots were generated using R, v. 4.5.1 (R Core Team (2025) with the EnhancedVolcano package(*129*). Molecular interaction and directionality of regulation was done using Search Tool for the Retrieval of Interacting Genes/Proteins (STRING) database analysis (version 12.0, medium confidence threshold score ≥ 0.4, disconnected nodes excluded)(*76*).

### Statistical analysis

All data were compared using GraphPad Prism (v 10.0.0). Normality was determined using the Shapiro-Wilk test. The differences between the two normally distributed groups were compared using a two-tailed unpaired Student’s t test. One-way analysis of variance (ANOVA) with Dunnett’s multiple comparisons test was used to compare multiple groups. The P values of <0.05, <0.01, and <0.001 were considered statistically significant differences and indicated as numbers in the charts (only statistically significant values are shown).

## Acknowledgements

Authors acknowledge the Biocenter Finland (BF), Tampere Imaging Facility (TIF), Tampere University Flow Cytometry Facility (TFCF), and Cornell Institute of Biotechnology’s BRC Imaging Core Facility (RRID:SCR_021741) for their services. Authors thank Assoc. Prof. Soile Nymark for the support and comments on the manuscript, Prof. Vesa Hytönen and Niklas Kähkönen for the production of Tn5 enzyme, Mervi Lindman for technical assistance with EM samples, the Electron Microscopy Unit (EMBI) of the Institute of Biotechnology, University of Helsinki (Helsinki, Finland) for EM sample preparation, Anja Hartewig for assistance in genetic analysis using IPA, and Prof. Pekka Katajisto for the transgenic mice and their genotyping.

## Funding

Research Council of Finland grant 308315 (TOI)

Research Council of Finland grant 314106 (TOI)

Research Council of Finland grant 332615 (EM)

Research Council of Finland grant 330896 (MVR)

Jane and Aatos Erkko Foundation (MVR)

European Union’s Horizon 2020 research and innovation programme, Compact Cell

Imaging Device (CoCID) under grant agreement No. 101017116 (MVR)

Finnish Cultural Foundation (JUM)

National Institutes of Health R01 GM137605 and R35 GM153257 (JL)

National Institute of Health R35 GM119617 (DEC)

The U.S. National Science Foundation award CMMI 2234888 (DEC)

## Author contributions

Conceptualization: TOI, EM

Data curation: TOI, EM, SH, JK

Formal analysis: TOI, EM, JK, SE, SK

Funding acquisition: TOI, EM, MVR, JUM, DEC, JL

Investigation: TOI, EM, AP, AR, SK, SA-MH, RD, MJTO, SH, JUM, AAV, PW, FA

Methodology: TOI, EM, JL, SK

Project administration: TOI, EM

Resources: TOI, EM, MN, MVR, KV, DEC, JL

Software: TOI, SE

Supervision: TOI, EM, KV, MVR, MN, DEC, JL

Validation: TOI, EM, JK, SH, JUM, RD, AAV, KV, JK, MN, DEC, JL

Visualization: TOI, EM, JK, SA-MH, SH, PW

Writing: EM, TOI, JK, SH, SA-MH, MJTO, RD, JUM, MVR, FA, JK

Writing – review & editing: TOI, EM, JK, AR, SH, SA-MH, MVR, AAV, JL, SK, RD, AP, MJTO, JUM, DEC, FA

## Data and materials availability

All data are available in the main text or the supplementary materials.

## References

1. C. G. Vasquez, A. C. Martin, Force Transmission in Epithelial Tissues. Dev. Dyn. Off. Publ. Am. Assoc. Anat. 245, 361–371 (2016).

2. V. Venturini, F. Pezzano, F. Català Castro, H.-M. Häkkinen, S. Jiménez-Delgado, M. Colomer-Rosell, M. Marro, Q. Tolosa-Ramon, S. Paz-López, M. A. Valverde, J. Weghuber, P. Loza-Alvarez, M. Krieg, S. Wieser, V. Ruprecht, The nucleus measures shape changes for cellular proprioception to control dynamic cell behavior. Science 370, eaba2644 (2020).

3. L. Schwartz, J. da Veiga Moreira, M. Jolicoeur, Physical forces modulate cell differentiation and proliferation processes. J. Cell. Mol. Med. 22, 738–745 (2018).

4. S. Blonski, J. Aureille, S. Badawi, D. Zaremba, L. Pernet, A. Grichine, S. Fraboulet, P. M. Korczyk, P. Recho, C. Guilluy, M. E. Dolega, Direction of epithelial folding defines impact of mechanical forces on epithelial state. Dev. Cell 56, 3222–3234.e6 (2021).

5. P. Strzyz, May the force be with you. Nat. Rev. Mol. Cell Biol. 17, 533–533 (2016).

6. C. Guilluy, L. D. Osborne, L. Van Landeghem, L. Sharek, R. Superfine, R. Garcia-Mata, K. Burridge, Isolated nuclei adapt to force and reveal a mechanotransduction pathway in the nucleus. Nat. Cell Biol. 16, 376–381 (2014).

7. G. V. Shivashankar, Mechanosignaling to the Cell Nucleus and Gene Regulation. Annu. Rev. Biophys. 40, 361–378 (2011).

8. S. Cho, J. Irianto, D. E. Discher, Mechanosensing by the nucleus: From pathways to scaling relationships. J. Cell Biol. 216, 305–315 (2017).

9. D. E. Ingber, Cellular mechanotransduction: Putting all the pieces together again. FASEB J. 20, 811–827 (2006).

10. C. S. Chen, Forces as regulators of cell adhesions. Nat. Rev. Mol. Cell Biol. 18, 715 (2017).

11. W. H. Goldmann, Mechanotransduction in cells. Cell Biol. Int. 36, 567–570 (2012).

12. J. Swift, D. E. Discher, The nuclear lamina is mechano-responsive to ECM elasticity in mature tissue. J. Cell Sci. 127, 3005–3015 (2014).

13. G. Chandramouly, P. C. Abad, D. W. Knowles, S. A. Lelièvre, The control of tissue architecture over nuclear organization is crucial for epithelial cell fate. J. Cell Sci. 120, 1596–1606 (2007).

14. F. Bertillot, Y. A. Miroshnikova, S. A. Wickström, SnapShot: Mechanotransduction in the nucleus. Cell 185, 3638–3638.e1 (2022).

15. T. O. Ihalainen, L. Aires, F. A. Herzog, R. Schwartlander, J. Moeller, V. Vogel, Differential basal-to-apical accessibility of lamin A/C epitopes in the nuclear lamina regulated by changes in cytoskeletal tension. Nat. Mater. 14, 1252–1261 (2015).

16. J. Paulsen, T. M. Liyakat Ali, M. Nekrasov, E. Delbarre, M.-O. Baudement, S. Kurscheid, D. Tremethick, P. Collas, Long-range interactions between topologically associating domains shape the four-dimensional genome during differentiation. Nat. Genet. 51, 835–843 (2019).

17. Q. Szabo, A. Donjon, I. Jerković, G. L. Papadopoulos, T. Cheutin, B. Bonev, E. P. Nora, B. G. Bruneau, F. Bantignies, G. Cavalli, Regulation of single-cell genome organization into TADs and chromatin nanodomains. Nat. Genet. 52, 1151–1157 (2020).

18. P. Niethammer, Components and Mechanisms of Nuclear Mechanotransduction. Annu. Rev. Cell Dev. Biol. 37, 233–256 (2021).

19. A. Tajik, Y. Zhang, F. Wei, J. Sun, Q. Jia, W. Zhou, R. Singh, N. Khanna, A. S. Belmont, N. Wang, Transcription upregulation via force-induced direct stretching of chromatin. Nat. Mater. 15, 1287–1296 (2016).

20. B. Seelbinder, S. Ghosh, A. G. Berman, S. E. Schneider, C. J. Goergen, S. Calve, C. P. Neu, Intra-Nuclear Tensile Strain Mediates Reorganization of Epigenetically Marked Chromatin During Cardiac Development and Disease. bioRxiv, 455600 (2019).

21. S.-J. Heo, B. D. Cosgrove, E. N. Dai, R. L. Mauck, Mechano-adaptation of the stem cell nucleus. Nucleus 9, 9–19 (2018).

22. Y. A. Miroshnikova, M. M. Nava, S. A. Wickström, Emerging roles of mechanical forces in chromatin regulation. J Cell Sci 130, 2243–2250 (2017).

23. B. van Steensel, A. S. Belmont, Lamina-Associated Domains: Links with Chromosome Architecture, Heterochromatin, and Gene Repression. Cell 169, 780–791 (2017).

24. E. Makhija, D. S. Jokhun, G. V. Shivashankar, Nuclear deformability and telomere dynamics are regulated by cell geometric constraints. Proc. Natl. Acad. Sci. 113, E32–E40 (2016).

25. C. H. Thomas, J. H. Collier, C. S. Sfeir, K. E. Healy, Engineering gene expression and protein synthesis by modulation of nuclear shape. Proc. Natl. Acad. Sci. 99, 1972–1977 (2002).

26. T. J. Kirby, J. Lammerding, Emerging views of the nucleus as a cellular mechanosensor. Nat. Cell Biol. 20, 373–381 (2018).

27. M. M. Nava, Y. A. Miroshnikova, L. C. Biggs, D. B. Whitefield, F. Metge, J. Boucas, H. Vihinen, E. Jokitalo, X. Li, J. M. García Arcos, B. Hoffmann, R. Merkel, C. M. Niessen, K. N. Dahl, S. A. Wickström, Heterochromatin-Driven Nuclear Softening Protects the Genome against Mechanical Stress-Induced Damage. Cell 181, 800–817.e22 (2020).

28. S. Hervé, Y. A. Miroshnikova, Biophysical determinants of nuclear shape and mechanics and their implications for genome integrity. Curr. Opin. Biomed. Eng. 30, 100521 (2024).

29. M. M. Drozdz, D. J. Vaux, Shared mechanisms in physiological and pathological nucleoplasmic reticulum formation. Nucleus 8, 34–45 (2017).

30. I. Singh, T. P. Lele, Nuclear morphological abnormalities in cancer – a search for unifying mechanisms. Results Probl. Cell Differ. 70, 443–467 (2022).

31. A. Malhas, C. Goulbourne, D. J. Vaux, The nucleoplasmic reticulum: form and function. Trends Cell Biol. 21, 362–373 (2011).

32. P. Marius, M. T. Guerra, M. H. Nathanson, B. E. Ehrlich, M. F. Leite, Calcium release from ryanodine receptors in the nucleoplasmic reticulum. Cell Calcium 39, 65–73 (2006).

33. A. J. Lomakin, C. J. Cattin, D. Cuvelier, Z. Alraies, M. Molina, G. P. F. Nader, N. Srivastava, P. J. Sáez, J. M. Garcia-Arcos, I. Y. Zhitnyak, A. Bhargava, M. K. Driscoll, E. S. Welf, R. Fiolka, R. J. Petrie, N. S. De Silva, J. M. González-Granado, N. Manel, A. M. Lennon-Duménil, D. J. Müller, M. Piel, The nucleus acts as a ruler tailoring cell responses to spatial constraints. Science 370, eaba2894 (2020).

34. B. Enyedi, M. Jelcic, P. Niethammer, The cell nucleus serves as a mechanotransducer of tissue damage-induced inflammation. Cell 165, 1160–1170 (2016).

35. M. Stiekema, F. Houben, F. Verheyen, M. Borgers, J. Menzel, M. Meschkat, M. A. M. J. van Zandvoort, F. C. S. Ramaekers, J. L. V. Broers, The Role of Lamins in the Nucleoplasmic Reticulum, a Pleiomorphic Organelle That Enhances Nucleo-Cytoplasmic Interplay. Front. Cell Dev. Biol. 10 (2022).

36. I. Schoen, L. Aires, J. Ries, V. Vogel, Nanoscale invaginations of the nuclear envelope: Shedding new light on wormholes with elusive function. Nucleus 8, 506–514 (2017).

37. I. Schoen, L. Aires, J. Ries, V. Vogel, Nanoscale invaginations of the nuclear envelope: Shedding new light on wormholes with elusive function. Nucleus 8, 506–514 (2017).

38. T.-C. Wang, C. R. Dollahon, S. Mishra, H. Patel, S. Abolghasemzade, I. Singh, V. Thomazy, D. G. Rosen, V. C. Sandulache, S. Chakraborty, T. P. Lele, Extreme wrinkling of the nuclear lamina is a morphological marker of cancer. *Npj Precis*. Oncol. 8, 1–12 (2024).

39. S. Schwertheim, S. Theurer, H. Jastrow, T. Herold, S. Ting, D. Westerwick, S. Bertram, C. M. Schaefer, J. Kälsch, H. A. Baba, K. W. Schmid, New insights into intranuclear inclusions in thyroid carcinoma: Association with autophagy and with BRAFV600E mutation. PLOS ONE 14, e0226199 (2019).

40. D. M. Jorgens, J. L. Inman, M. Wojcik, C. Robertson, H. Palsdottir, W.-T. Tsai, H. Huang, A. Bruni-Cardoso, C. S. López, M. J. Bissell, K. Xu, M. Auer, Deep nuclear invaginations are linked to cytoskeletal filaments – integrated bioimaging of epithelial cells in 3D culture. J. Cell Sci. 130, 177–189 (2017).

41. B. D. Cosgrove, C. Loebel, T. P. Driscoll, T. K. Tsinman, E. N. Dai, S.-J. Heo, N. A. Dyment, J. A. Burdick, R. L. Mauck, Nuclear envelope wrinkling predicts mesenchymal progenitor cell mechano-response in 2D and 3D microenvironments. Biomaterials 270, 120662 (2021).

42. N. Johnson, M. Krebs, R. Boudreau, G. Giorgi, M. LeGros, C. Larabell, Actin-filled nuclear invaginations indicate degree of cell de-differentiation. Differentiation 71, 414–424 (2003).

43. D. M. Jorgens, J. L. Inman, M. Wojcik, C. Robertson, H. Palsdottir, W.-T. Tsai, H. Huang, A. Bruni-Cardoso, C. S. López, M. J. Bissell, K. Xu, M. Auer, Deep nuclear invaginations are linked to cytoskeletal filaments – integrated bioimaging of epithelial cells in 3D culture. J. Cell Sci. 130, 177–189 (2017).

44. A. Mukherjee, J. E. Ron, H. T. Hu, T. Nishimura, K. Hanawa-Suetsugu, B. Behkam, Y. Mimori-Kiyosue, N. S. Gov, S. Suetsugu, A. S. Nain, Actin Filaments Couple the Protrusive Tips to the Nucleus through the I-BAR Domain Protein IRSp53 during the Migration of Cells on 1D Fibers. Adv. Sci. 10, 2207368 (2023).

45. G. Rappa, M. F. Santos, T. M. Green, J. Karbanová, J. Hassler, Y. Bai, S. H. Barsky, D. Corbeil, A. Lorico, Nuclear transport of cancer extracellular vesicle-derived biomaterials through nuclear envelope invagination-associated late endosomes. Oncotarget 8, 14443–14461 (2017).

46. J. Geng, Z. Kang, Q. Sun, M. Zhang, P. Wang, Y. Li, J. Li, B. Su, Q. Wei, Microtubule Assists Actomyosin to Regulate Cell Nuclear Mechanics and Chromatin Accessibility. Research, doi: 10.34133/research.0054 (2023).

47. H. Cheng, C. P. Leblond, Origin, differentiation and renewal of the four main epithelial cell types in the mouse small intestine. V. Unitarian Theory of the origin of the four epithelial cell types. Am. J. Anat. 141, 537–561 (1974).

48. S. B. Khatau, C. M. Hale, P. J. Stewart-Hutchinson, M. S. Patel, C. L. Stewart, P. C. Searson, D. Hodzic, D. Wirtz, A perinuclear actin cap regulates nuclear shape. Proc. Natl. Acad. Sci. 106, 19017–19022 (2009).

49. J.-K. Kim, A. Louhghalam, G. Lee, B. W. Schafer, D. Wirtz, D.-H. Kim, Nuclear lamin A/C harnesses the perinuclear apical actin cables to protect nuclear morphology. Nat. Commun. 8, 2123 (2017).

50. S. Biedzinski, G. Agsu, B. Vianay, M. Delord, L. Blanchoin, J. Larghero, L. Faivre, M. Théry, S. Brunet, Microtubules control nuclear shape and gene expression during early stages of hematopoietic differentiation. EMBO J. 39, e103957 (2020).

51. F. Paonessa, L. D. Evans, R. Solanki, D. Larrieu, S. Wray, J. Hardy, S. P. Jackson, F. J. Livesey, Microtubules Deform the Nuclear Membrane and Disrupt Nucleocytoplasmic Transport in Tau-Mediated Frontotemporal Dementia. Cell Rep. 26, 582–593.e5 (2019).

52. A. J. Sarria, J. G. Lieber, S. K. Nordeen, R. M. Evans, The presence or absence of a vimentin-type intermediate filament network affects the shape of the nucleus in human SW-13 cells. J. Cell Sci. 107, 1593–1607 (1994).

53. T. Abe, K. Takano, A. Suzuki, Y. Shimada, M. Inagaki, N. Sato, T. Obinata, T. Endo, Myocyte differentiation generates nuclear invaginations traversed by myofibrils associating with sarcomeric protein mRNAs. J. Cell Sci. 117, 6523–6534 (2004).

54. F. Alisafaei, D. S. Jokhun, G. V. Shivashankar, V. B. Shenoy, Regulation of nuclear architecture, mechanics, and nucleocytoplasmic shuttling of epigenetic factors by cell geometric constraints. Proc. Natl. Acad. Sci. 116, 13200–13209 (2019).

55. P. T. Arsenovic, I. Ramachandran, K. Bathula, R. Zhu, J. D. Narang, N. A. Noll, C. A. Lemmon, G. G. Gundersen, D. E. Conway, Nesprin-2G, a Component of the Nuclear LINC Complex, Is Subject to Myosin-Dependent Tension. Biophys. J. 110, 34–43 (2016).

56. P. M. Davidson, J. Lammerding, Broken nuclei – lamins, nuclear mechanics, and disease. Trends Cell Biol. 24, 247–256 (2014).

57. L. K. Srivastava, Z. Ju, A. Ghagre, A. J. Ehrlicher, Spatial distribution of lamin A/C determines nuclear stiffness and stress-mediated deformation. J. Cell Sci. 134, jcs248559 (2021).

58. B. E. Danielsson, B. G. Abraham, E. Mäntylä, J. I. Cabe, C. R. Mayer, A. Rekonen, F. Ek, D. E. Conway, T. O. Ihalainen, Nuclear lamina strain states revealed by intermolecular force biosensor. bioRxiv, 2022.03.07.483300 (2022).

59. P. M. Davidson, J. Sliz, P. Isermann, C. Denais, J. Lammerding, Design of a microfluidic device to quantify dynamic intra-nuclear deformation during cell migration through confining environments. Integr. Biol. Quant. Biosci. Nano Macro 7, 1534–1546 (2015).

60. P. M. Davidson, C. Denais, M. C. Bakshi, J. Lammerding, Nuclear deformability constitutes a rate-limiting step during cell migration in 3-D environments. Cell. Mol. Bioeng. 7, 293–306 (2014).

61. M. Wallace, G. R. Fedorchak, R. Agrawal, R. M. Gilbert, J. Patel, S. Park, M. Paszek, J. Lammerding, The lamin A/C Ig-fold undergoes cell density-dependent changes that alter epitope binding. Nucleus 14, 2180206 (2023).

62. E. Carley, R. M. Stewart, A. Zieman, I. Jalilian, D. E. King, A. Zubek, S. Lin, V. Horsley, M. C. King, The LINC complex transmits integrin-dependent tension to the nuclear lamina and represses epidermal differentiation. eLife 10, e58541 (2021).

63. Y. Kalukula, A. D. Stephens, J. Lammerding, S. Gabriele, Mechanics and functional consequences of nuclear deformations. Nat. Rev. Mol. Cell Biol. 23, 583–602 (2022).

64. X. Chen, Y. Shen, W. Draper, J. D. Buenrostro, U. Litzenburger, S. W. Cho, A. T. Satpathy, A. C. Carter, R. P. Ghosh, A. East-Seletsky, J. A. Doudna, W. J. Greenleaf, J. T. Liphardt, H. Y. Chang, ATAC-see reveals the accessible genome by transposase-mediated imaging and sequencing. Nat. Methods 13, 1013–1020 (2016).

65. F. Greil, C. Moorman, B. van Steensel, DamID: mapping of in vivo protein-genome interactions using tethered DNA adenine methyltransferase. Methods Enzymol. 410, 342–359 (2006).

66. A. Puliafito, L. Hufnagel, P. Neveu, S. Streichan, A. Sigal, D. K. Fygenson, B. I. Shraiman, Collective and single cell behavior in epithelial contact inhibition. Proc. Natl. Acad. Sci. U. S. A. 109, 739–744 (2012).

67. M. A. Nieto, Epithelial Plasticity: A Common Theme in Embryonic and Cancer Cells. Science 342, 1234850 (2013).

68. S. Li, E. R. Gerrard, D. F. Balkovetz, Evidence for ERK1/2 phosphorylation controlling contact inhibition of proliferation in Madin-Darby canine kidney epithelial cells. Am. J. Physiol. Cell Physiol. 287, C432–439 (2004).

69. Y. Matsubayashi, M. Ebisuya, S. Honjoh, E. Nishida, ERK Activation Propagates in Epithelial Cell Sheets and Regulates Their Migration during Wound Healing. Curr. Biol. 14, 731–735 (2004).

70. A. Liberzon, C. Birger, H. Thorvaldsdóttir, M. Ghandi, J. P. Mesirov, P. Tamayo, The Molecular Signatures Database (MSigDB) hallmark gene set collection. Cell Syst. 1, 417–425 (2015).

71. Ingenuity Pathway Analysis, Bioinformatics Software. QIAGEN Digital Insights. https://digitalinsights.qiagen.com/products-overview/discovery-insights-portfolio/analysis-and-visualization/qiagen-ipa/ (accessed 10 March 2026).

72. A. D. Rouillard, G. W. Gundersen, N. F. Fernandez, Z. Wang, C. D. Monteiro, M. G. McDermott, A. Ma’ayan, The harmonizome: a collection of processed datasets gathered to serve and mine knowledge about genes and proteins. Database 2016, baw100 (2016).

73. I. Diamant, D. J. B. Clarke, J. E. Evangelista, N. Lingam, A. Ma’ayan, Harmonizome 3.0: integrated knowledge about genes and proteins from diverse multi-omics resources. Nucleic Acids Res. 53, D1016–D1028 (2025).

74. A. Krämer, J. Green, J. Pollard, S. Tugendreich, Causal analysis approaches in Ingenuity Pathway Analysis. Bioinforma. Oxf. Engl. 30, 523–530 (2014).

75. T. Boulding, F. Wu, R. McCuaig, J. Dunn, C. R. Sutton, K. Hardy, W. Tu, A. Bullman, D. Yip, J. E. Dahlstrom, S. Rao, Differential Roles for DUSP Family Members in Epithelial-to-Mesenchymal Transition and Cancer Stem Cell Regulation in Breast Cancer. PLOS ONE 11, e0148065 (2016).

76. D. Szklarczyk, R. Kirsch, M. Koutrouli, K. Nastou, F. Mehryary, R. Hachilif, A. L. Gable, T. Fang, N. T. Doncheva, S. Pyysalo, P. Bork, L. J. Jensen, C. von Mering, The STRING database in 2023: protein-protein association networks and functional enrichment analyses for any sequenced genome of interest. Nucleic Acids Res. 51, D638–D646 (2023).

77. E. Shaulian, M. Karin, AP-1 as a regulator of cell life and death. Nat. Cell Biol. 4, E131–136 (2002).

78. M. Olea-Flores, M. D. Zuñiga-Eulogio, M. A. Mendoza-Catalán, H. A. Rodríguez-Ruiz, E. Castañeda-Saucedo, C. Ortuño-Pineda, T. Padilla-Benavides, N. Navarro-Tito, Extracellular-Signal Regulated Kinase: A Central Molecule Driving Epithelial–Mesenchymal Transition in Cancer. Int. J. Mol. Sci. 20, 2885 (2019).

79. A. Gutierrez-Hartmann, D. L. Duval, A. P. Bradford, ETS transcription factors in endocrine systems. Trends Endocrinol. Metab. 18, 150–158 (2007).

80. W. Zhang, H. T. Liu, MAPK signal pathways in the regulation of cell proliferation in mammalian cells. Cell Res. 12, 9–18 (2002).

81. A. Pastuła, R. Lundmark, Induction of Epithelial-mesenchymal Transition in MDCK II Cells. Bio-Protoc. 11, e3903 (2021).

82. M. Davies, M. Robinson, E. Smith, S. Huntley, S. Prime, I. Paterson, Induction of an epithelial to mesenchymal transition in human immortal and malignant keratinocytes by TGF-beta1 involves MAPK, Smad and AP-1 signalling pathways. J. Cell. Biochem. 95, 918–931 (2005).

83. J. M. González, A. Navarro-Puche, B. Casar, P. Crespo, V. Andrés, Fast regulation of AP-1 activity through interaction of lamin A/C, ERK1/2, and c-Fos at the nuclear envelope. J. Cell Biol. 183, 653–666 (2008).

84. E. Mäntylä, T. O. Ihalainen, Brick Strex: a robust device built of LEGO bricks for mechanical manipulation of cells. Sci. Rep. 11, 18520 (2021).

85. A. Liberzon, C. Birger, H. Thorvaldsdóttir, M. Ghandi, J. P. Mesirov, P. Tamayo, The Molecular Signatures Database (MSigDB) hallmark gene set collection. Cell Syst. 1, 417–425 (2015).

86. M. F. Favata, K. Y. Horiuchi, E. J. Manos, A. J. Daulerio, D. A. Stradley, W. S. Feeser, D. E. Van Dyk, W. J. Pitts, R. A. Earl, F. Hobbs, R. A. Copeland, R. L. Magolda, P. A. Scherle, J. M. Trzaskos, Identification of a novel inhibitor of mitogen-activated protein kinase kinase. J. Biol. Chem. 273, 18623–18632 (1998).

87. S. Y. Liu, K. Ikegami, Nuclear lamin phosphorylation: an emerging role in gene regulation and pathogenesis of laminopathies. Nucleus 11, 299–314 (2020).

88. J. Koester, Y. A. Miroshnikova, S. Ghatak, C. A. Chacón-Martínez, J. Morgner, X. Li, I. Atanassov, J. Altmüller, D. E. Birk, M. Koch, W. Bloch, M. Bartusel, C. M. Niessen, A. Rada-Iglesias, S. A. Wickström, Niche stiffening compromises hair follicle stem cell potential during ageing by reducing bivalent promoter accessibility. Nat. Cell Biol. 23, 771–781 (2021).

89. F. Broders-Bondon, T. H. Nguyen Ho-Bouldoires, M.-E. Fernandez-Sanchez, E. Farge, Mechanotransduction in tumor progression: The dark side of the force. J. Cell Biol. 217, 1571–1587 (2018).

90. H. Jayatilaka, F. G. Umanzor, V. Shah, T. Meirson, G. Russo, B. Starich, P. Tyle, J. S. H. Lee, S. Khatau, H. Gil-Henn, D. Wirtz, H. Jayatilaka, F. G. Umanzor, V. Shah, T. Meirson, G. Russo, B. Starich, P. Tyle, J. S. H. Lee, S. Khatau, H. Gil-Henn, D. Wirtz, H. Jayatilaka, F. G. Umanzor, V. Shah, T. Meirson, G. Russo, B. Starich, P. Tyle, J. S. H. Lee, S. Khatau, H. Gil-Henn, D. Wirtz, Tumor cell density regulates matrix metalloproteinases for enhanced migration. Oncotarget 9, 32556–32569 (2018).

91. V. von Manstein, B. Groner, Tumor cell resistance against targeted therapeutics: the density of cultured glioma tumor cells enhances Stat3 activity and offers protection against the tyrosine kinase inhibitor canertinib. MedChemComm 8, 96–102 (2017).

92. B. Coste, J. Mathur, M. Schmidt, T. J. Earley, S. Ranade, M. J. Petrus, A. E. Dubin, A. Patapoutian, Piezo1 and Piezo2 are essential components of distinct mechanically activated cation channels. Science 330, 55–60 (2010).

93. C. D. Cox, N. Bavi, B. Martinac, Biophysical Principles of Ion-Channel-Mediated Mechanosensory Transduction. Cell Rep. 29, 1–12 (2019).

94. J. J. Northey, L. Przybyla, V. M. Weaver, Tissue Force Programs Cell Fate and Tumor Aggression. Cancer Discov. 7, 1224 LP–1237 (2017).

95. S. Dupont, S. A. Wickström, Mechanical regulation of chromatin and transcription. Nat. Rev. Genet. 23, 624–643 (2022).

96. A. Karoutas, A. Akhtar, Functional mechanisms and abnormalities of the nuclear lamina. Nat. Cell Biol. 23, 116–126 (2021).

97. C. S. Xu, K. J. Hayworth, Z. Lu, P. Grob, A. M. Hassan, J. G. García-Cerdán, K. K. Niyogi, E. Nogales, R. J. Weinberg, H. F. Hess, Enhanced FIB-SEM systems for large-volume 3D imaging. eLife 6, e25916 (2017).

98. L. Heinrich, D. Bennett, D. Ackerman, W. Park, J. Bogovic, N. Eckstein, A. Petruncio, J. Clements, S. Pang, C. S. Xu, J. Funke, W. Korff, H. F. Hess, J. Lippincott-Schwartz, S. Saalfeld, A. V. Weigel, Whole-cell organelle segmentation in volume electron microscopy. Nature 599, 141–146 (2021).

99. C. M. Denais, R. M. Gilbert, P. Isermann, A. L. McGregor, M. te Lindert, B. Weigelin, P. M. Davidson, P. Friedl, K. Wolf, J. Lammerding, Nuclear envelope rupture and repair during cancer cell migration. Science 352, 353–8 (2016).

100. P. Scaffidi, T. Misteli, Lamin A-Dependent Nuclear Defects in Human Aging. Science 312, 1059–1063 (2006).

101. B. McKee, S. Abolghasemzade, T.-C. Wang, K. Harsh, S. Kaur, R. Blanchard, K. B. Menon, M. Mohajeri, R. B. Dickinson, T. P. Lele, Excess surface area of the nuclear lamina enables unhindered cell migration through constrictions. Sci. Adv. 11, eads6573 (2025).

102. A. Bermudez, Z. D. Latham, A. J. Ma, D. Bi, J. K. Hu, N. Y. C. Lin, Regulation of chromatin modifications through coordination of nucleus size and epithelial cell morphology heterogeneity. Commun. Biol. 8, 269 (2025).

103. L. Guelen, L. Pagie, E. Brasset, W. Meuleman, M. B. Faza, W. Talhout, B. H. Eussen, A. de Klein, L. Wessels, W. de Laat, B. van Steensel, Domain organization of human chromosomes revealed by mapping of nuclear lamina interactions. Nature 453, 948–951 (2008).

104. A. K. Efremov, L. Hovan, J. Yan, Nucleus size and its effect on nucleosome stability in living cells. Biophys. J. 121, 4189–4204 (2022).

105. M. P. Creyghton, A. W. Cheng, G. G. Welstead, T. Kooistra, B. W. Carey, E. J. Steine, J. Hanna, M. A. Lodato, G. M. Frampton, P. A. Sharp, L. A. Boyer, R. A. Young, R. Jaenisch, Histone H3K27ac separates active from poised enhancers and predicts developmental state. Proc. Natl. Acad. Sci. 107, 21931–21936 (2010).

106. A. M. Mendonsa, T.-Y. Na, B. M. Gumbiner, E-cadherin in contact inhibition and cancer. Oncogene 37, 4769–4780 (2018).

107. H. Liu, L. Yuan, L. Baldi, T. R. Sornapudi, G. V. Shivashankar, Compressive Forces Induce Epigenetic Activation of Aged Human Dermal Fibroblasts Through ERK Signaling Pathway. Aging Cell 24, e70035 (2025).

108. J. Schindelin, I. Arganda-Carreras, E. Frise, V. Kaynig, M. Longair, T. Pietzsch, S. Preibisch, C. Rueden, S. Saalfeld, B. Schmid, J.-Y. Tinevez, D. J. White, V. Hartenstein, K. Eliceiri, P. Tomancak, A. Cardona, Fiji: an open-source platform for biological-image analysis. Nat. Methods 9, 676–682 (2012).

109. P. W. Tillberg, F. Chen, K. D. Piatkevich, Y. Zhao, C.-C. Yu, B. P. English, L. Gao, A. Martorell, H.-J. Suk, F. Yoshida, E. M. DeGennaro, D. H. Roossien, G. Gong, U. Seneviratne, S. R. Tannenbaum, R. Desimone, D. Cai, E. S. Boyden, Protein-retention expansion microscopy of cells and tissues labeled using standard fluorescent proteins and antibodies. Nat. Biotechnol. 34, 987–992 (2016).

110. J. Moeller, A. K. Denisin, J. Y. Sim, R. E. Wilson, A. J. S. Ribeiro, B. L. Pruitt, Controlling cell shape on hydrogels using lift-off protein patterning. PloS One 13, e0189901 (2018).

111. S. Picelli, Å. K. Björklund, B. Reinius, S. Sagasser, G. Winberg, R. Sandberg, Tn5 transposase and tagmentation procedures for massively scaled sequencing projects. Genome Res. 24, 2033–2040 (2014).

112. J. D. Buenrostro, P. G. Giresi, L. C. Zaba, H. Y. Chang, W. J. Greenleaf, Transposition of native chromatin for fast and sensitive epigenomic profiling of open chromatin, DNA-binding proteins and nucleosome position. Nat. Methods 10, 1213–1218 (2013).

113. Babraham Bioinformatics - FastQC A Quality Control tool for High Throughput Sequence Data. https://www.bioinformatics.babraham.ac.uk/projects/fastqc/.

114. Y. Zhang, T. Liu, C. A. Meyer, J. Eeckhoute, D. S. Johnson, B. E. Bernstein, C. Nusbaum, R. M. Myers, M. Brown, W. Li, X. S. Liu, Model-based Analysis of ChIP-Seq (MACS). Genome Biol. 9, R137 (2008).

115. Babraham Bioinformatics - Trim Galore! https://www.bioinformatics.babraham.ac.uk/projects/trim_galore/.

116. B. Langmead, S. L. Salzberg, Fast gapped-read alignment with Bowtie 2. Nat. Methods 9, 357–359 (2012).

117. M. R. Corces, J. M. Granja, S. Shams, B. H. Louie, J. A. Seoane, W. Zhou, T. C. Silva, C. Groeneveld, C. K. Wong, S. W. Cho, A. T. Satpathy, M. R. Mumbach, K. A. Hoadley, A. G. Robertson, N. C. Sheffield, I. Felau, M. A. A. Castro, B. P. Berman, L. M. Staudt, J. C. Zenklusen, P. W. Laird, C. Curtis, The Cancer Genome Atlas Analysis Network, W. J. Greenleaf, H. Y. Chang, The chromatin accessibility landscape of primary human cancers. Science 362, eaav1898 (2018).

118. A. T. L. Lun, G K. Smyth, csaw: a Bioconductor package for differential binding analysis of ChIP-seq data using sliding windows. Nucleic Acids Res. 44, e45 (2016).

119. J. J. Reske, M. R. Wilson, R. L. Chandler, ATAC-seq normalization method can significantly affect differential accessibility analysis and interpretation. Epigenetics Chromatin 13, 22 (2020).

120. G. Yu, L.-G. Wang, Q.-Y. He, ChIPseeker: an R/Bioconductor package for ChIP peak annotation, comparison and visualization. Bioinformatics 31, 2382–2383 (2015).

121. S. Heinz, C. Benner, N. Spann, E. Bertolino, Y. C. Lin, P. Laslo, J. X. Cheng, C. Murre, H. Singh, C. K. Glass, Simple combinations of lineage-determining transcription factors prime cis-regulatory elements required for macrophage and B cell identities. Mol. Cell 38, 576–589 (2010).

122. S. Anders, W. Huber, Differential expression analysis for sequence count data. Genome Biol. 11, R106 (2010).

123. V. K. Mootha, C. M. Lindgren, K.-F. Eriksson, A. Subramanian, S. Sihag, J. Lehar, P. Puigserver, E. Carlsson, M. Ridderstråle, E. Laurila, N. Houstis, M. J. Daly, N. Patterson, J. P. Mesirov, T. R. Golub, P. Tamayo, B. Spiegelman, E. S. Lander, J. N. Hirschhorn, D. Altshuler, L. C. Groop, PGC-1α-responsive genes involved in oxidative phosphorylation are coordinately downregulated in human diabetes. Nat. Genet. 34, 267–273 (2003).

124. A. Subramanian, P. Tamayo, V. K. Mootha, S. Mukherjee, B. L. Ebert, M. A. Gillette, A. Paulovich, S. L. Pomeroy, T. R. Golub, E. S. Lander, J. P. Mesirov, Gene set enrichment analysis: a knowledge-based approach for interpreting genome-wide expression profiles. Proc. Natl. Acad. Sci. U. S. A. 102, 15545–15550 (2005).

125. V. K. Mootha, C. M. Lindgren, K.-F. Eriksson, A. Subramanian, S. Sihag, J. Lehar, P. Puigserver, E. Carlsson, M. Ridderstråle, E. Laurila, N. Houstis, M. J. Daly, N. Patterson, J. P. Mesirov, T. R. Golub, P. Tamayo, B. Spiegelman, E. S. Lander, J. N. Hirschhorn, D. Altshuler, L. C. Groop, PGC-1α-responsive genes involved in oxidative phosphorylation are coordinately downregulated in human diabetes. Nat. Genet. 34, 267–273 (2003).

126. H. Heberle, G. V. Meirelles, F. R. da Silva, G. P. Telles, R. Minghim, InteractiVenn: a web-based tool for the analysis of sets through Venn diagrams. BMC Bioinformatics 16, 169 (2015).

127. A. Krämer, J. Green, J. Pollard Jr, S. Tugendreich, Causal analysis approaches in Ingenuity Pathway Analysis. Bioinformatics 30, 523–530 (2014).

128. QIAGEN IPA Interpret, Bioinformatics Software. QIAGEN Digital Insights. https://digitalinsights.qiagen.com/products-overview/discovery-insights-portfolio/analysis-and-visualization/qiagen-ipa/qiagen-ipa-interpret/ (accessed 10 March 2026).

129. K. Blighe, J. Andrews, S. Rana, E. Turkes, B. Ostendorf, A. Grioni, M. Lewis, EnhancedVolcano: Publication-ready volcano plots with enhanced colouring and labeling, Bioconductor (2025). 10.18129/B9.bioc.EnhancedVolcano.

130. N. Barker, J. H. van Es, J. Kuipers, P. Kujala, M. van den Born, M. Cozijnsen, A. Haegebarth, J. Korving, H. Begthel, P. J. Peters, H. Clevers, Identification of stem cells in small intestine and colon by marker gene Lgr5. Nature 449, 1003–1007 (2007).

131. T. Sato, H. Clevers, Growing Self-Organizing Mini-Guts from a Single Intestinal Stem Cell: Mechanism and Applications. Science 340, 1190–1194 (2013).

132. Y. Kasagi, P. M. Chandramouleeswaran, K. A. Whelan, K. Tanaka, V. Giroux, M. Sharma, J. Wang, A. J. Benitez, M. DeMarshall, J. W. Tobias, K. E. Hamilton, G. W. Falk, J. M. Spergel, A. J. Klein-Szanto, A. K. Rustgi, A. B. Muir, H. Nakagawa, The Esophageal Organoid System Reveals Functional Interplay Between Notch and Cytokines in Reactive Epithelial Changes. Cell. Mol. Gastroenterol. Hepatol. 5, 333–352 (2018).

