## Supplemental Materials for "Deep Invaginations of Nuclear Envelope Coordinate Spatial Organization of Chromatin in Epithelium"

**Contents:**

Figs. S1 to S5
Tables S1 to S5
Movies S1 to S3 (.avi)
Supplementary Materials and Methods for Fig. S2

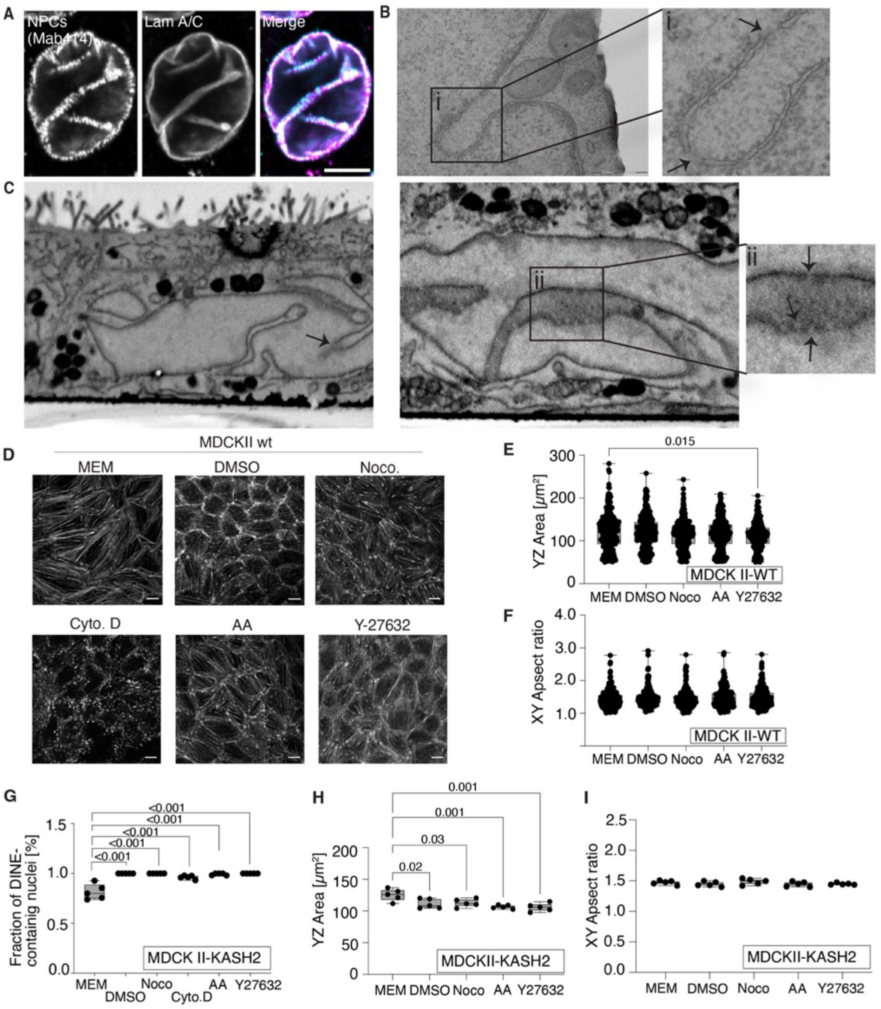


**Fig. S1. Characterization of DINEs and role of cytoskeleton and LINC-mediated coupling in DINE maintenance. (A)** Representative LSCM images showing nuclear pore complexes (NPCs) in lamin A/C immunostained peripheral NE and DINEs. Scale bar, 5 µm. **(B)** TEM image showing NPCs in DINEs deep within the nucleus indicating potentially functional NE. **(C)** Volume EM images indicating DINEs traversing through the nucleus and the presence of NPCs in the DINEs within the nucleus. Arrows indicate single NPCS. **(D)** LSCM images of phalloidin-stained basal actin cytoskeleton in non-treated control (Modified Eagle´s Medium, MEM), and cytoskeleton disruptive drug-treated (DMSO, Nocodazole (Noco.), Cytochalasin D (Cyto. D), acrylamide (AA), and ROCK-inhibitor Y-27632) MDCK II wt cells. Scale bars, 10 µm. Box- and whiskers plot showing quantification of **(E)** nuclear YZ area (µm$^2$), and **(F)** nuclear XY aspect ratio in drug-treated WT cells (n=20 fields with 246, 260, 277, 247, and 271 cells, respectively, three independent biological replicates), and **(G)** mean fraction (%) of DINE-containing nuclei per field of view (n=5 fields with 223, 275, 278, 266, and 271 cells, respectively, three independent biological replicates, One-way ANOVA with Dunnett´s multiple comparison test, $p<0.05$ indicates statistical significance), **(H)** nuclear YZ area (µm$^2$), and **(I)** nuclear XY aspect ratio in non-treated or drug-treated MDCK II-KASH2 cells (n= one independent biological replicate, One-way ANOVA with Dunnett´s multiple comparison test, $p<0.05$ indicates statistical significance). Box-and-whisker plots represent the 25$^{th}$–75$^{th}$ percentiles as boxes, the median as a line within the box, and whiskers indicating the minimum and maximum values.

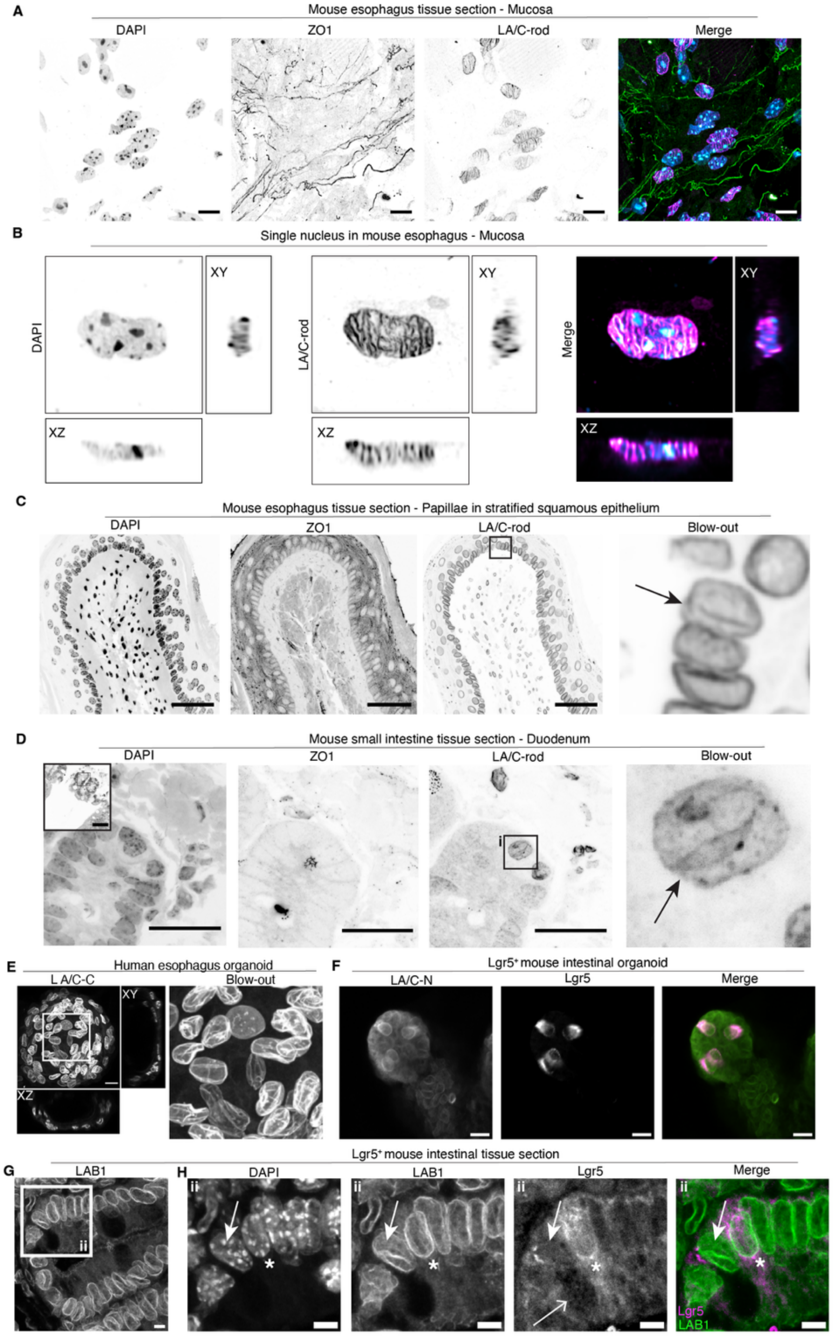


**Fig. S2. DINEs as an innate property of epithelial nuclei *in vitro* and *in vivo*. (A)** LSCM image of DAPI, ZO1 and LA/C-rod immunostained mouse esophagus tissue section (mucosa). Scale bars, 10 µm. **(B)** Planar and orthogonal views of a single DINE-containing DAPI and LA/C-rod -stained nucleus in mouse esophagus mucosa. LSCM images of DAPI, ZO1 and LA/C-rod immunostained **(C)** mouse esophagus tissue sections showing papillae in stratified squamous epithelium including a DINE-containing nucleus, and **(D)** mouse small intestine (duodenum) tissue sections (black arrows indicate a DINE-containing nucleus). Scale bars, 25 µm. **(E)** LA/C-rod -stained human esophagus 3D organoid with DINE-containing nuclei. Scale bar, 20 µm. **(F)** Mouse intestinal organoid expressing Lgr5-EGFP positive (Lgr5$^{+}$) intestinal stem cells with smooth nuclear lamina avoid of DINEs detected with an antibody targeting LA/C-N. Scale bars, 20 µm. **(G)** Lgr5-EGFP positive mouse intestinal tissue section and **(H)** blow-out of area ii indicated in (G) showing DINE-containing nucleus (bolded white arrow) in a secretory cell identified by the presence of cytoplasmic granules (thin white arrow) next to a Lgr5$^{+}$ stem cell (white asterisk) immunostained with LAB1, DAPI, and Turbo-GFP detecting Lgr5-EGFP. Scale bars, 5 µm.

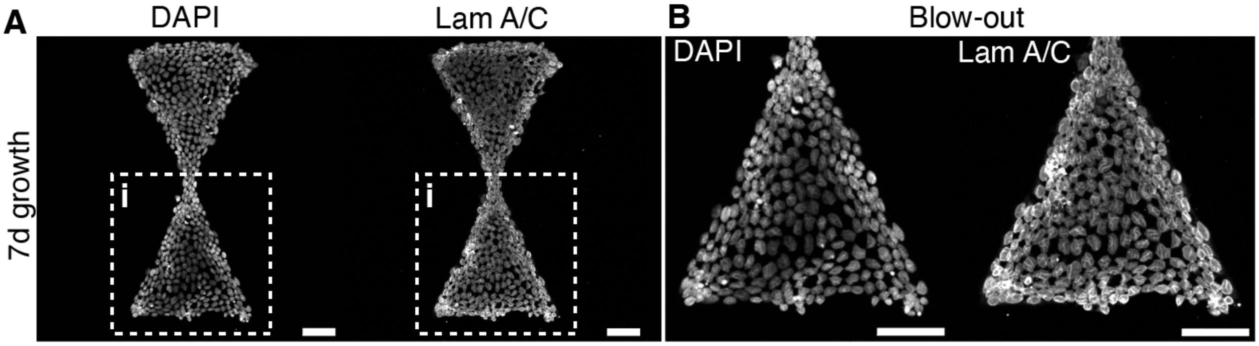


**Fig. S3. Influence of constricted 2D growth surface and 3D migration on DINEs. (A)** LSCM images and **(B)** blow out of area i shown in (A) of LA-CB cells grown for 7 d on PLL-PEG-TRITC-stained hourglass-shaped cell adhesive islands, showing increased cellular growth density and DINE-containing nuclei on the micropattern edges and waist area. Chromatin and lamin A/C were detected with DAPI and LA/C-rod antibodies, respectively. Scale bars, 50 µm.

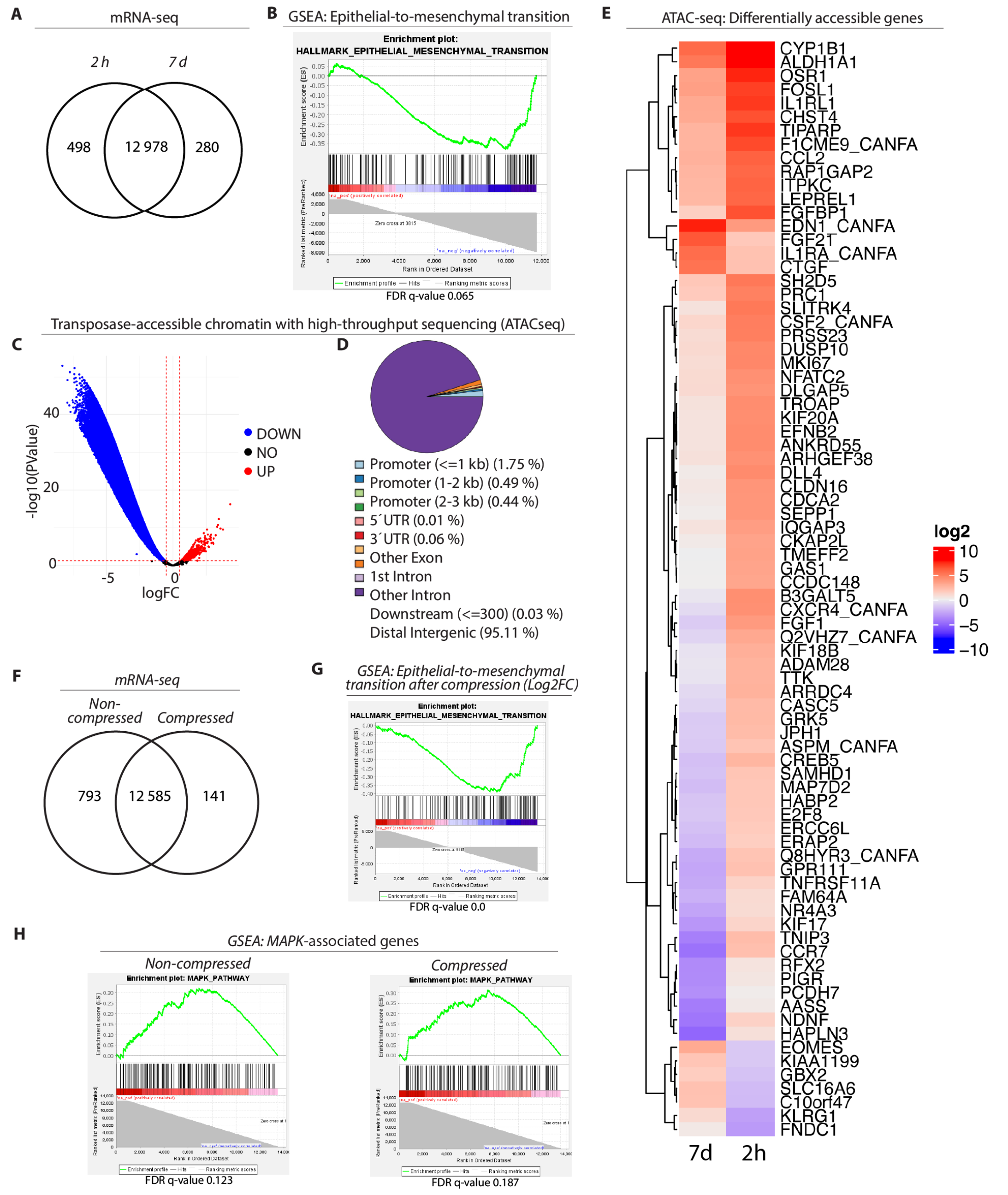


**Fig. S4. Analysis of unique gene expression, EMT-status, transcriptionally accessible chromatin, and intracellular localization of pERK. (A)** Venn-diagram showing mRNA-seq -derived unique and commonly expressed genes in epithelial cells grown for 7 d or in cells grown for 7 d and reseeded for 2 h. **(B)** Gene Set Enrichment Analysis (GSEA) showing mostly negative expression of EMT-associated genes at 7 d (FDR value 0.065). **(C)** Volcano plot showing transcriptionally accessible and inaccessible chromatin detected with assay for transposase-accessible chromatin using sequencing (ATAC-seq) indicating that most chromatin in 7 d DINE-containing nuclei was transcriptionally inaccessible in comparison to those grown for 2 h after reseeding from a 7 d -grown epithelium. **(D)** Pie chart indicating that over 95 % of the accessible chromatin regions were mapping mainly to distal intergenic sequences. **(E)** Heatmap indicating genes with significantly increased (blue) or decreased accessibility between 7 d and 2 h grown samples following ATAC-seq. **(F)** Venn-diagram showing mRNA-seq -derived unique and commonly expressed genes in epithelial cells before and after the lateral compression. GSEA indicating **(G)** positive and negative enrichment of EMT-associated genes following the compression, and **(H)** positive and negative enrichment of MAPK-associated genes before (non-compressed, left) and after (compressed, right) compression is shown.

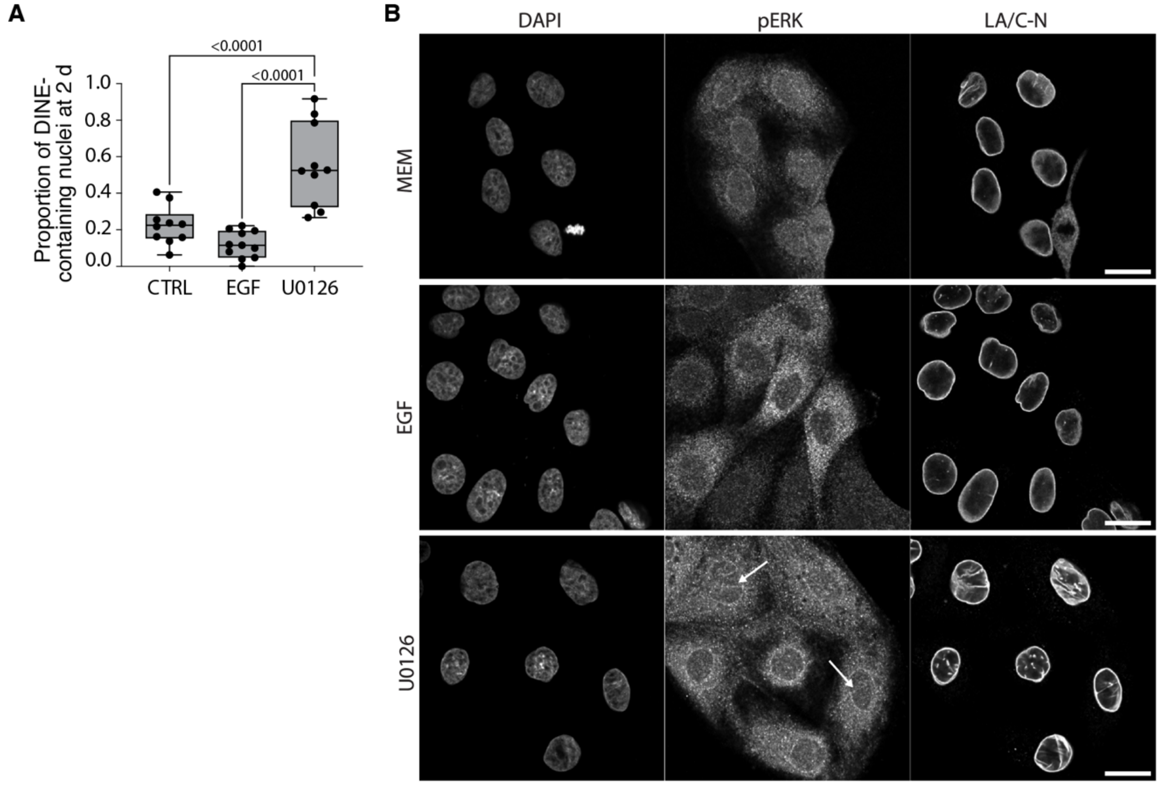


**Fig. S5. Quantification of DINE-containing nuclei at 2 d, intracellular distribution of active phosphorylated ERK1/2 following pharmacological induction and suppression of epithelial-to-mesenchymal transition (EMT). (A)** Quantification showing DINE-containing nuclei in developing monolayer plated without (CTRL) or in the presence of drugs (ERK1/2 activator epidermal growth factor EGF or MEK1/2 inhibitor U0126) and grown for 2 d followed by fixing and LA/C-rod and DAPI immunostaining prior quantifications. One-way ANOVA with Dunnett´s multiple comparison test, $p<0.05$ indicates statistical significance), n = two independent biological replicates. Box-and-whisker plots represent the $25^{th}$–$75^{th}$ percentiles as boxes, the median as a line within the box, and whiskers indicating the minimum and maximum values. **(B)** LSCM images showing nucleo-cytoplasmic presence of active phosphorylated ERK1/2 (pERK) in LA/C-N and phospho-ERK1/2 (pERK) immunostained cells grown for 2 d without drugs (in medium, MEM, upper panels), increased cytoplasmic expression and spotted nuclear presence following EMT activation with EGF (100 ng/mL, middle panels), and cytoplasmic and NE-associated presence of pERK following EMT inhibition with U0126 (50 µM, lower panels). White arrowheads indicate localization of pERK at the NE and in the DINEs, n= two independent biological replicates. Scale bars, 20 µm.

**Table S1. Effects of cytoskeleton disruptive drugs on nuclear morphology.** Nuclear area (xy, μm$^2$), yz aspect ratio and circularity of 7-d grown wild-type (wt) and dominant negative KASH2 cells without drugs or following a 2 h treatment with DMSO, nocodazole, acrylamide and ROCK-inhibitor Y27632. Values presented as mean ± standard deviation (SD).

| | Treatment | Nuclear area [μm$^2$] | YZ Aspect ratio | Circularity |
|---|---|---|---|---|
| WT | MEM | 122 ± 44 | 1.43 ± 0.27 | 0.79 ± 0.06 |
| | DMSO | 121 ± 39 | 1.47 ± 0.28 | 0.79 ± 0.06 |
| | Nocodazole | 114 ± 34 | 1.44 ± 0.27 | 0.79 ± 0.06 |
| | Acrylamide | 117 ± 36 | 1.48 ± 0.29 | 0.79 ± 0.06 |
| | Y27632 | 112 ± 31 | 1.48 ± 0.29 | 0.79 ± 0.06 |
| KASH2 | MEM | 125 ± 9.5 | 1.47 ± 0.03 | 0.80 ± 0.01 |
| | DMSO | 111 ± 7.9 | 1.45 ± 0.04 | 0.80 ± 0.01 |
| | Nocodazole | 104 ± 6.4 | 1.49 ± 0.05 | 0.79 ± 0.01 |
| | Acrylamide | 106 ± 2.9 | 1.45 ± 0.04 | 0.79 ± 0.01 |
| | Y27632 | 106 ± 6.5 | 1.45 ± 0.02 | 0.79 ± 0.01 |

**Table S2. Ingenuity pathway analysis (IPA).** Excel file indicating IPA analysis results ((-log(p-value)) > 1.3 and |z-score| >2) with significant differential expressed genes (DEGs) acquired from mRNA-seq done in mature epithelium grown for 7 d and in laterally compressed epithelial monolayer after 2 h recovery. The log(p-value), ratio, z-score and name of the molecules are shown.

**Table S3. IPA MAPK-pathway analysis of mature and compressed epithelium.** Excel file listing the name, Log2FoldChange (Log2FC), and up- or downregulation of significant differential expressed genes acquired from mRNA-seq done in 7 d grown mature epithelium containing DINEs, and compressed epithelial monolayer containing DINEs.

**Table S4. Search Tool for the Retrieval of Interacting Genes/Proteins (STRING) database analysis of mature epithelium grown for 7 d.** Excel file indicating STRING analysis -derived interactions of molecules in nodes (neighborhood on chromosome, gene fusion, phylogenetic cooccurrence, homology, coexpression, experimentally determined interaction, database annotated, automated_textmining, combined score), functional annotations, and cluster analysis (kmeans clusters, kmeans cluster descriptions) (supports Fig. 6).

**Table S5. ATAC-seq MAPK-associated transcription factor (TF) motif analysis of mature epithelium grown for 7 d.** Excel file containing the list of MAPK-associated TFs used in the ATAC-seq motif analysis.

**Movie S1. Deep invaginations of the nuclear envelope (DINEs).** Surface rendering of an expansion microscopy (ExM)-LSCM -imaged single epithelial cell nucleus immunostained with LA/C-C (green) and LAB1 (red) showing DINEs protruding from the basal toward the apical nuclear surface. Scale bar, 5 µm.

**Movie S2. DINEs in human esophagus organoid.** 3D rendering of an LA/C-rod -immunostained human esophagus organoid showing DINEs in a 3D tissue scaffold. Scale bar, 20 µm.

**Movie S3. Confined 3D migration of a DINE-containing epithelial cell.** Time lapse imaging of LA-CB -expressing MDCK II cells indicating unfolding of a DINE before squeezing of the nucleus through a thin ($2 \times 5$ $mm^2$) constriction within a microfluidic 3D migration device. Imaging 6 fph. Scale bar, 10 mm.

### Supplementary Materials and Methods

#### Materials and Methods for Supplemental Figure S2.

##### Animals

Four- to nine-month-old Lgr5-EGFP-IRES-creERT2 mice (JAX stock #008875, The Jackson Laboratory, ME, USA) (130) on a mixed C57BL/6JRccHsd and C57BL/6N background were used in the study. The transgenic mice were genotyped with PCR using the following primers: 5´ CTGCTCTCTGCTCCCAGTCT´3 (common primer), 5´ ATACCCCATCCCTTTTGAGC´3 (wild type primer) and 5´GAACTTCAGGGTCAGCTTGC´3 (GFP primer). The mice were maintained in the pathogen-free preclinical facility of Tampere University (Tampere, Finland) under the permit ESAVI/33102/2024 and according to the Finnish Act on the Protection of Animals Used for Scientific or Educational Purposes (497/2013) and the EU Directive (2010/63/EU) guidelines. Mice were housed in individually ventilated cages at regulated temperature (22 °C) and humidity (45-65 %) with a 12-hour light/dark cycle. Chow and water were provided ad libitum.

##### Intestinal Organoid Culture

3D organoid culture was done as originally described by the protocol of Sato et al. 2009 (131). Intestinal segments were opened longitudinally, washed thoroughly with ice-cold PBS to remove luminal contents. Intestinal piece was aspirated and dispensed through 10 ml serological pipet to remove villus. For detaching crypts, 10 mmol/L EDTA was administered for 30 min incubation in +4°C on gentle shaking. Chelation-based dissociation 1:30 ratio of fetal bovine serum (FBS) was used to enrich for intact crypts by aspirating and dispensing through 5ml serological pipet. Crypt-containing fractions were collected, washed with cold 1xPBS and pelleted. Crypts were embedded in 35 µl domes in 50% Matrigel (#256231; Corning). Cultures were maintained at 37 °C and 5% $CO_2$. Crypts were cultured in an optimal medium consisting of advanced Dulbecco's modified Eagle medium/F12 (#12634010; Thermo Fisher Scientific, Waltham, MA, USA) supplemented with, Glutamax (2 mmol/L, # 35050061; Thermo Fisher Scientific), HEPES (10 mmol/L, #15630-080; Sigma-Aldrich, St. Louis, MO), penicillin-streptomycin (100 U/mL, #11659990; Sigma-Aldrich), B-27 supplement (#17504-044; Thermo Fisher Scientific), N-2 supplement (#17502-001; Thermo Fisher Scientific), N-acetylcysteine (1 mmol/L, # A9165; Sigma-Aldrich), recombinant murine epidermal growth factor (50 ng/mL, # PMG8043, lot 2135273; Gibco, Waltham, MA), recombinant murine Noggin (100 ng/mL, #250-38; PeproTech), and recombinant human R-spondin-1 (1 µg/mL, #120-38; Peprotech). Medium was refreshed every 2–3 days, and for culture establishment only first 2 days a ROCK inhibitor Y-27632 (#10005583; Cayman Chemicals) was included. Organoids were passaged by recovering domes, mechanically dissociating into smaller fragments/crypt units, re-embedding in fresh matrix, and returning to growth medium.

##### Human esophageal organoid culture

Biopsies from normal esophageal mucosal epithelium were collected during upper endoscopy. Primary cell isolation and esophageal organoid culture were carried out following the protocol described by Kasagi et al. 2018 (132), with the modifications outlined below. Esophageal fragments were washed with 10 mL of ice-cold 1X PBS and collected by centrifugation at 400 x g for 5 minutes at 4°C. The supernatant was discarded, and the washed esophageal fragments were incubated with 1 mL of Collagenase/Dispase (1 mg/mL; #10269638001, Roche) at 37°C for 45 minutes with gentle shaking and vigorous pipetting every 10 minutes to facilitate dissociation. An additional 1 mL of 0.05 % sterile Trypsin-EDTA (#T2601, Sigma-Aldrich) was added to the fragments and incubated at 37°C for 10 minutes with gentle shaking. The dissociation enzymes were neutralized by adding 1 mL of fetal bovine serum (#F1283, Sigma-Aldrich) and 7 mL of ice-cold 1X PBS. The entire content was filtered through a 40 µm cell strainer (#CLS431750, Corning), followed by a centrifugation step. The supernatant was discarded, and an additional wash with ice-cold 1X PBS was performed. Dissociated esophageal epithelial cells were resuspended in undiluted Matrigel (#256231, Corning) and seeded in 35 µL domes per well in a 24-well multidish (#142475, Thermo Scientific). The plate was then incubated at 37°C for 5 minutes to allow complete Matrigel polymerization. The organoids were cultured in Keratinocyte-SFM (#17005042, Gibco) supplemented with bovine pituitary extract (30 µg/mL; #13028-014, Gibco), human recombinant epidermal growth factor (0.2 ng/mL; #10450-013, Gibco), penicillin-streptomycin (100 U/mL; #11659990, Sigma-Aldrich), and CaCl2 (0.6mM; #J63122, Thermo Scientific Chemicals). The ROCK inhibitor Y-27632 (#10005583, Cayman Chemicals) was added to the medium during the first two days to facilitate culture establishment. Cultures were maintained at 37°C and 5% CO2, and the medium was refreshed every 48 hours. Esophageal organoids were passaged on day 12. Matrigel domes were harvested in 10 mL of ice-cold 1X PBS and collected by gentle centrifugation. After discarding the supernatant, the sedimented organoids were incubated in 1 mL of TrypLE Express Enzyme (1M; #12604013, Gibco) at 37°C for 20 minutes with gentle shaking and vigorous pipetting every 5 minutes. TrypLE was neutralized by adding 9 mL of ice-cold 1X PBS, followed by filtration and centrifugation. The resulting single cells were re-embedded in 35 µL Matrigel domes and cultured in supplemented KSFM growth medium. For staining, dissociated single cells were embedded in Matrigel and plated on an 8-well Corning Falcon culture slides (#CLS354118, Corning) at a density of $1x10^3$ cells/well.

**Immunohistochemistry of tissue sections**

Tissues were fixed in 10 % formalin for 24 h. Mouse esophagus (mucosa and papillae in stratified squamous epithelium) and small intestine (duodenum) samples were cut into 400 or 500µm thick paraffin sections using a microtome (Leica SM 2010R, Leica Biosystems, Illinois, United States) and attached to glass slides with a 1h incubation at 60 °C. To remove paraffin, the tissues were first treated with xylene or UltraClear™ (#3905.2500PE, J.T. Baker, United Kingdom) for 5 min, 4 min, 3 min and 1-1.5 min followed with absolute EtOH for 2 x 4 min, 96 % EtOH for 2 x 3 min and 70 % EtOH for 3 min and rinsed in distilled $H_2O$ (dH2O). Antigen retrieval for the esophagus was done by boiling in 10 mM sodium citrate (pH 6) or for the duodenal samples in 10 mM Tris- 1 mM EDTA (pH 9) and samples cooled down. Next, the samples were washed twice in TBS with 0.05% Tween-20 (TBST) and blocked by incubating them in TBS with 10 % fetal bovine serum (FBS) and 5 % BSA (esophagus) or with 1% BSA (duodenum) in a humid chamber for 1 h at 37 °C. For the antibody staining, primary Abs (mouse monoclonal anti-ZO1 antibody (ZO1-1A12, Invitrogen, Thermo Fisher Scientific, MA, USA), rabbit recombinant monoclonal anti-LA/C-rod ([EP4520, ab133256, Abcam, Cambridge, UK), GFP-Booster Alexa Fluor 488 (#gb2AF488, Chromotek, Planegg-Martinsried, Germany), mouse monoclonal anti-LA/C-N ([E1], sc-376248, Santa Cruz Biotechnology, TE, USA) and polyclonal anti-lamin B1 (#ab16048, Abcam) were diluted in TBS with 1 % BSA according to manufacturer instructions, the samples washed twice with TBST for 5 min, and the antibodies incubated with the tissue section in a humid chamber o/n at 4 °C. After primary Ab incubation, samples were washed twice in TBST for 5 min. The secondary Abs Alexa Fluor 568–conjugated goat anti-rabbit IgG (#A11011, Thermo Fisher Scientific), Alexa Fluor 488–conjugated goat anti-mouse IgG (#A-11001, Thermo Fisher Scientific) were added according to manufacturer instructions in TBST or in in-house made Mouse on Mouse -diluent for 1 h in RT, and finally washed twice in TBST for 5 min. Samples were mounted in Prolong Diamond Antifade with DAPI and covered with a Zeiss High Performance (18 x 18 mm) or a Menzel-Gläser (24x60 mm) glass. Samples were cured o/n in RT, in dark, and stored in 4 °C prior to imaging.
